# Mechanically-induced Septin Networks Protect Nuclear Integrity

**DOI:** 10.64898/2026.01.20.700414

**Authors:** Margaret E. Utgaard, Alexia Caillier, Shreya Chandrasekar, Joseph J. Tidei, Harikrushnan Balasubramanian, Rachel M. Lee, Owen Puls, Satya Khuon, Jesse Aaron, Teng-Leong Chew, Jordan R. Beach, Patrick W. Oakes

## Abstract

The cytoskeleton is a key mediator of mechanical interactions in cells, but specific contributions of septins remains unclear. Septins preferentially localize with a subset of actin stress fibers positioned under the nucleus, where they are situated between the membrane and stress fibers. Removing the nucleus from the cell results in the loss of these subnuclear septin-decorated stress fibers. Surprisingly, however, their formation can be rescued using a large glass bead in place of the nucleus. Similarly, applying a compressive force to the cell via confinement, whether externally or through internally generated actomyosin forces, results in increased septin accumulation in regions where the nucleus engages the cell cortex. Finally, loss of septin filaments via knockdown of SEPT7 increases the likelihood of nuclear membrane rupture during confinement. Together these data suggest that septins act as a dynamic mechanosensitive protective mechanism to buffer mechanical forces on the nucleus.

## Introduction

Physiological environments are crowded. Cells are surrounded by other cells and dense extracellular matrix, whose mechanical properties change throughout development and disease. When coupled with the internally generated forces from the actomyosin cytoskeleton, these confined architectures apply significant stress and strain on cells, even when cells appear stationary [1]. On account of its size and stiffness, the nucleus is particularly sensitive to mechanical forces and is often the limiting factor for navigating through confined spaces [2–5]. While deformation of the nucleus can contribute to downstream mechanotransduction [6, 7], excessive deformation can cause DNA damage, loss of nuclear integrity, and ultimately apoptosis [3, 8]. Cells therefore must employ defenses to buffer these mechanical forces.

The cytoskeleton is largely responsible for regulating both a cell’s morphology and its mechanical properties. As dynamic networks of filaments, the organization and architecture of the cytoskeleton is responsive to a variety of biochemical and mechanical cues. Generally, myosin motors pulling on the actin cytoskeleton create contractile networks that keep cells globally under tension, while microtubules and intermediate filaments aid in resisting compressive forces [9]. Each of these cytoskeletal networks have unique mechanical properties that can change both spatially and temporally in response to changing environmental conditions [9, 10]. Complicating the matter, they do not operate independently and interactions with one another can further modulate their behavior and mechanics [11–14].

The septin cytoskeleton is most well-known for its role in cell division, but also prominently localizes to subnuclear stress fibers and other cytoskeletal structures during interphase, where they likely scaffold signaling molecules [15–19]. Septins largely form heteromeric palindromic subunits, comprised of monomers from three or four different families of septin genes [18, 20]. Like other cytoskeletal elements, septin subunits dynamically assemble into filaments, and can bundle and crosslink into larger networks [15, 20]. Septins can also directly bundle and stabilize both actin and microtubules, providing a clear connection between these networks [21–23]. Intriguingly, septins are found at many sites where there are large applied forces, such as the contractile ring, sperm annulus, cilia base, and stress fibers [24–26]. Despite these localizations and interactions, however, septins direct role in mechanotransduction remains ill-defined.

Here we set out to investigate why septins preferentially localize to the subset of actin stress fibers located around the nucleus. We first created an endogenously-tagged SEPT2 fibroblast cell line to visualize septins in live-imaging under various perturbations. We observe that perinuclear septin localization is dynamic, moving with the nucleus during migration, and that it is present in both 3D and on substrates of physiological stiffness. Using a combination of super-resolution microscopy approaches we find that septins are typically positioned below actin stress fibers, closer to the membrane. Enucleating cells results in the loss of perinuclear septin fibers that can be partially recovered by introducing large glass beads into the cytoplasm, indicating a role for mechanics in septin formation. Applying a compressive force to the cells via confinement results in increased septin accumulation under the nucleus, and this behavior is mimicked as cells squeeze themselves through constrictions in a microfluidic device. In each case, septins accumulate in regions where the nucleus is pressed against the cortex. Lastly, we knock down SEPT7 and show that without these septin networks, nuclei are more susceptible to deformation under strain and ultimately more likely to rupture in confinement. Together these findings suggest that septins have a novel role in mechanotransduction, specifically by playing a dynamic protective role buffering mechanical forces on the nucleus.

## Results

### Septins form a dynamic network around the nucleus

As perturbation of individual septin genes has been previously reported to affect expression levels of other septin monomers that comprise the heteromeric subunit [15], we used CRISPR/Cas9 to endogenously-tag SEPT2 with a C-terminal HaloTag to visualize septins in a fibroblast cell line (SEPT2-Halo; Fig. S1A). SEPT2-Halo perfectly colocalized with antibody staining of SEPT2, SEPT7, and SEPT9 indicating that it can successfully incorporate into filaments, while Western blots revealed no impact on protein levels (Fig. S1B,C). Similar to previous reports [15–17], these stainings reveal septin accumulation along portions of the cortex, and a robust subnuclear septin network that aligns with actin stress fibers (Figs. 1A, S1B). This nuclear network is not exclusive to the ventral surface, as enrichment also occurs along actin stress fibers above the nucleus on the dorsal surface (Fig. 1A). Comparable accumulations are seen around the nucleus when cells are plated on elastic substrates or embedded in 3D collagen matrices, indicating that this localization occurs regardless of the substrate stiffness and architecture (Fig. S2).

**Figure 1.**
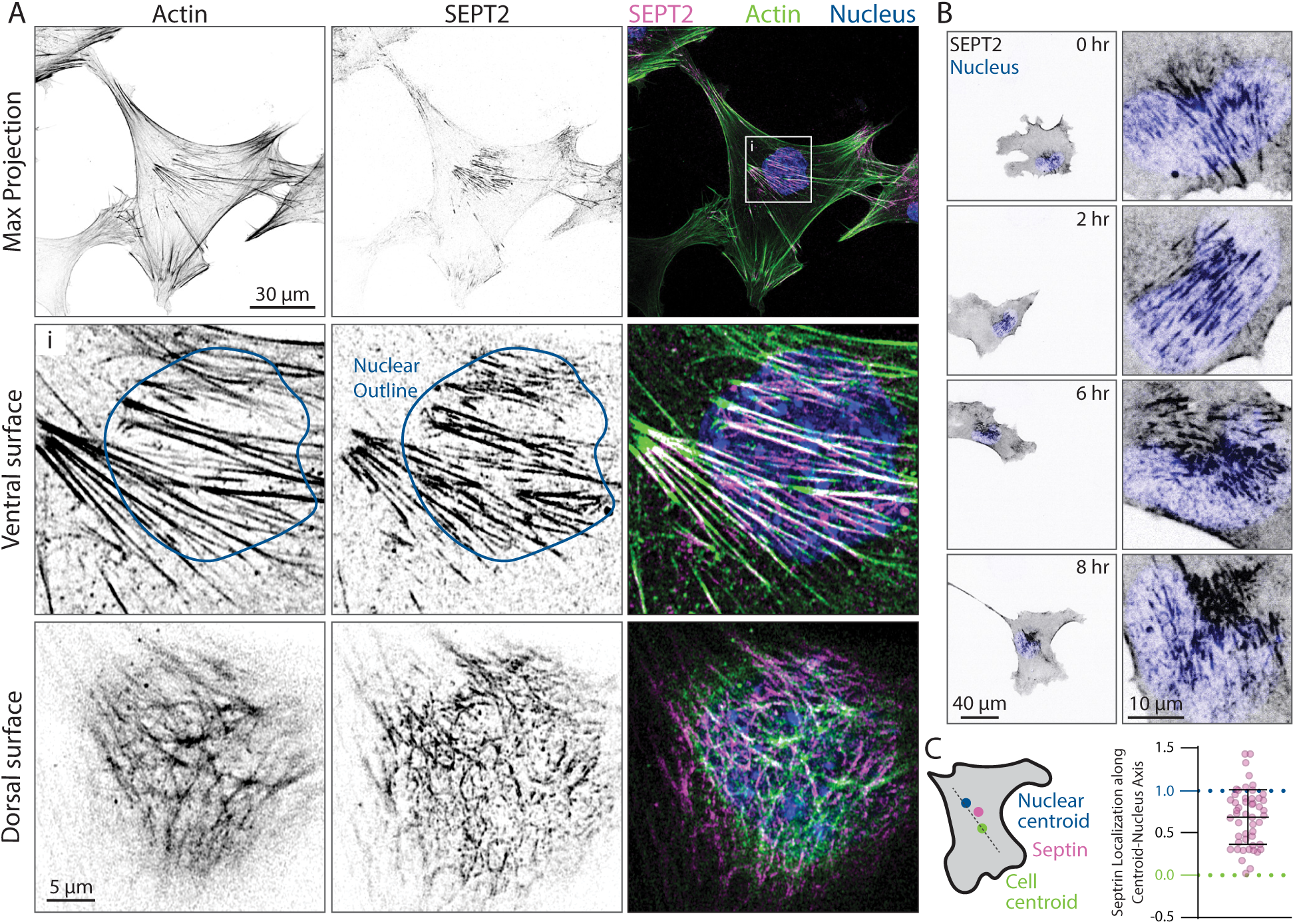
Septins coincide and move with the nucleus. **A)** Fibroblasts were fixed, immunolabeled with SEPT2 antibody (magenta), and stained for actin (phalloidin; green) and nuclei (DAPI; blue). The top row is a max projection of the whole cell, while the middle and bottom rows are single confocal slices of cropped regions (white box) above and below the nucleus. **B)** SEPT2-Halo cells stained with Hoechst nuclear dye were imaged for 12 hrs (Movie M1). Select time-points are shown of the whole cell in the left column and cropped regions near the nucleus in the right column. **C)** The center of intensity of the ventromedial septin network along the axis running from the cell centroid, defined as 0, to the nuclear centroid, defined as 1, was identified in SEPT2-Halo cells stained for nuclei (DAPI). Positive values indicate a septin preference towards the nucleus and negative numbers indicate a septin preference away from the nucleus. Data was collected from 48 cells (gray dots) from 3 independent experiments, with error bars indicate mean ± SD.

During migration this perinuclear septin network dynamically reorganizes as the cell changes morphology (Fig. 1B and Movie M1). Specifically, as the cell moves the septin network reorganizes to remain primarily under the nucleus, although extensions beyond the nuclear periphery occur. To quantify this spatial correlation, we measured the cell centroid, nuclear centroid, and center of intensity of the prominent perinuclear septin network (Fig. 1C). For all cells, the center of the septin network was positively biased towards the nucleus relative to the cell centroid, supporting a model that the ventral septin network and the nucleus are spatially coupled (Fig. 1C).

### Septin networks form between the actin and plasma membrane

Given the clear relationship between septins, actin, and the nucleus, we sought to gain a better understanding of the organization of septins and actin in these structures. We performed three-dimensional structured illumination microscopy (SIM) and found that the prominent septin filaments formed both below and above, but not along the sides of the nucleus (Fig. 2A). Further underscoring this point, the subnuclear actin and septin fibers often extend beyond the boundary of the nucleus along the cortex (Figs. 1A, 2A). These observations suggest that these septin and actin networks are situated at the cortex and not the nuclear envelope.

**Figure 2.**
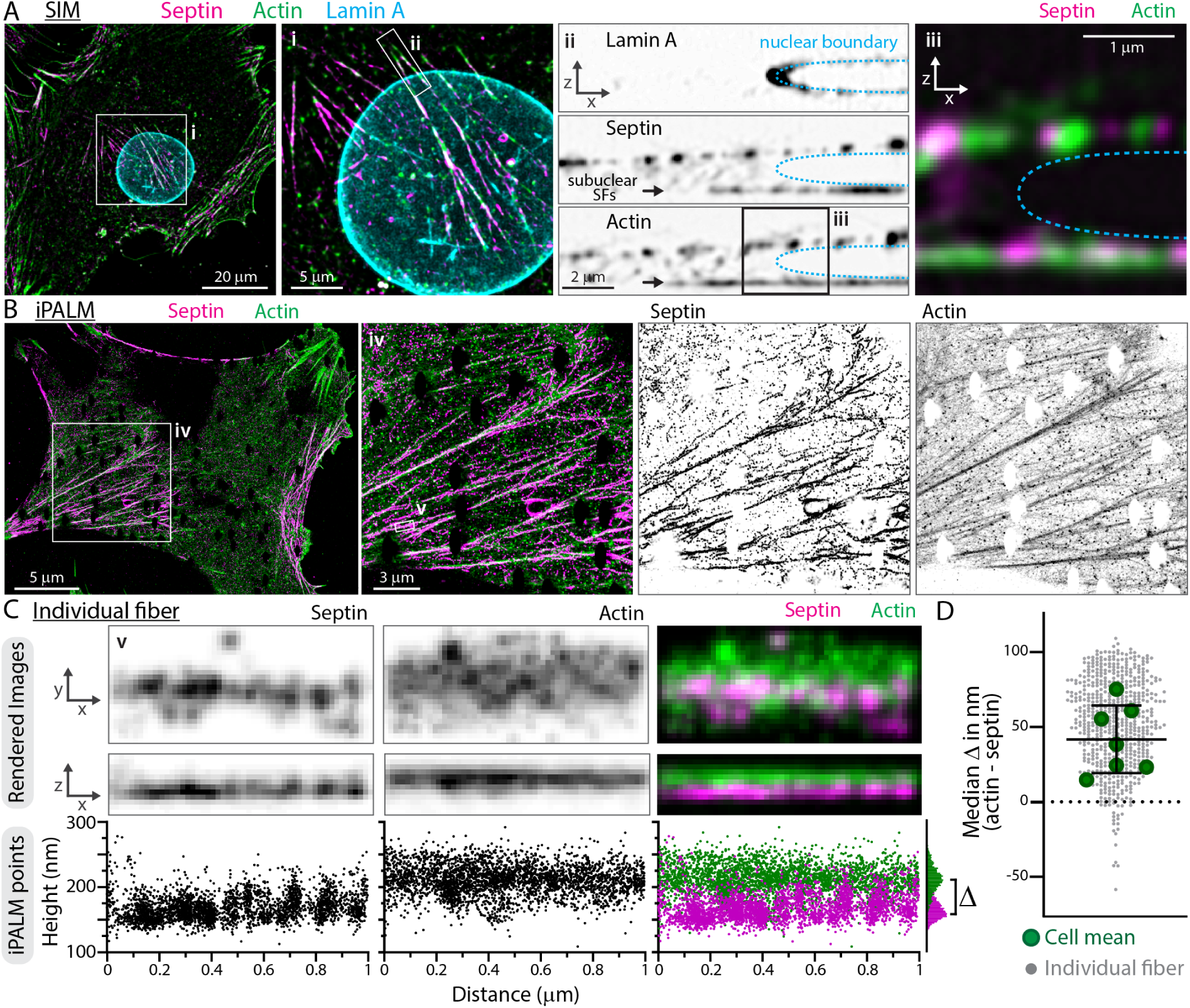
Septins are positioned along the cortex below actin. **A)** SIM imaging of SEPT2-Halo, actin (FastAct_X) and mEmerald-LaminA. Whole-cell and nuclear zoom are mean projections of short ventral z-stacks for actin and septin, and a mean projection of the entire z-stack for laminA. Individual inverted grayscale and merged orthogonal images in (ii and iii) are mean projections with nuclear boundary indicated by dashed cyan line. **B)** Whole-cell and subnuclear regions of rendered iPALM images. **C)** Rendered individual fiber (top), rendered orthogonal view (middle), and orthogonal plot of iPALM points from the same fiber (bottom). **D)** The difference (Δ) between the median actin and median septin height was calculated for 599 fibers from 7 cells, and plotted for each fiber (gray dots) and cell means (green dots) with error bars indicating mean ± SD of cell means.

To explore this hypothesis further, we used interference photoactivation localization microscopy (iPALM) to further resolve the spatial organization of the actin and septin networks under the nucleus. iPALM imaging, which provides nanometer scale lateral and axial resolution, reveals that despite having overlapping X-Y localizations, septins are consistently positioned below actin filaments (Fig. 2B,C). To quantify this separation, we isolated individual subnuclear fibers and measured the difference in the median position of the septin and actin localization events to be approximately 45 nm (Fig. 2D). While this distribution was broad, it was consistently positive. This distinct separation of the septin and actin network suggests septins could potentially have a unique role in cortical and nuclear mechanics.

### Externally applied forces induce septin network formation under the nucleus

To investigate the role of mechanical signaling in stimulating septin network formation, we used a previously developed dynamic confinement system to observe septin assembly in response to external compression [27]. We chose a confinement height of 5 microns to apply compressive pressure to the nucleus without causing catastrophic mechanical damage. Upon compression, septin intensity accumulates under the nucleus (Fig. 3A,B, Movie M2). While the compression occurs rapidly (< 1 min), the changes in septin distribution occur over 10s of minutes (Fig. 3A), indicating that the subnuclear signal increase is not due to pushing pre-assembled septin networks down into the ventral imaging plane. Upon release of compression, the septin accumulation under the nucleus returns to pre-compression levels (Fig. 3A,B, Movie M2). Imaging of actin, using EGFP-FTractin, and myosin, using EGFP-non-muscle myosin 2A (EGFP-NM2A) reveals similar increases in these cytoskeletal components under the nucleus (Fig. 3A,B, Movie M2). These data demonstrate nuclear compression into the cortex via extracellularly applied force elicits a septin assembly response.

**Figure 3.**
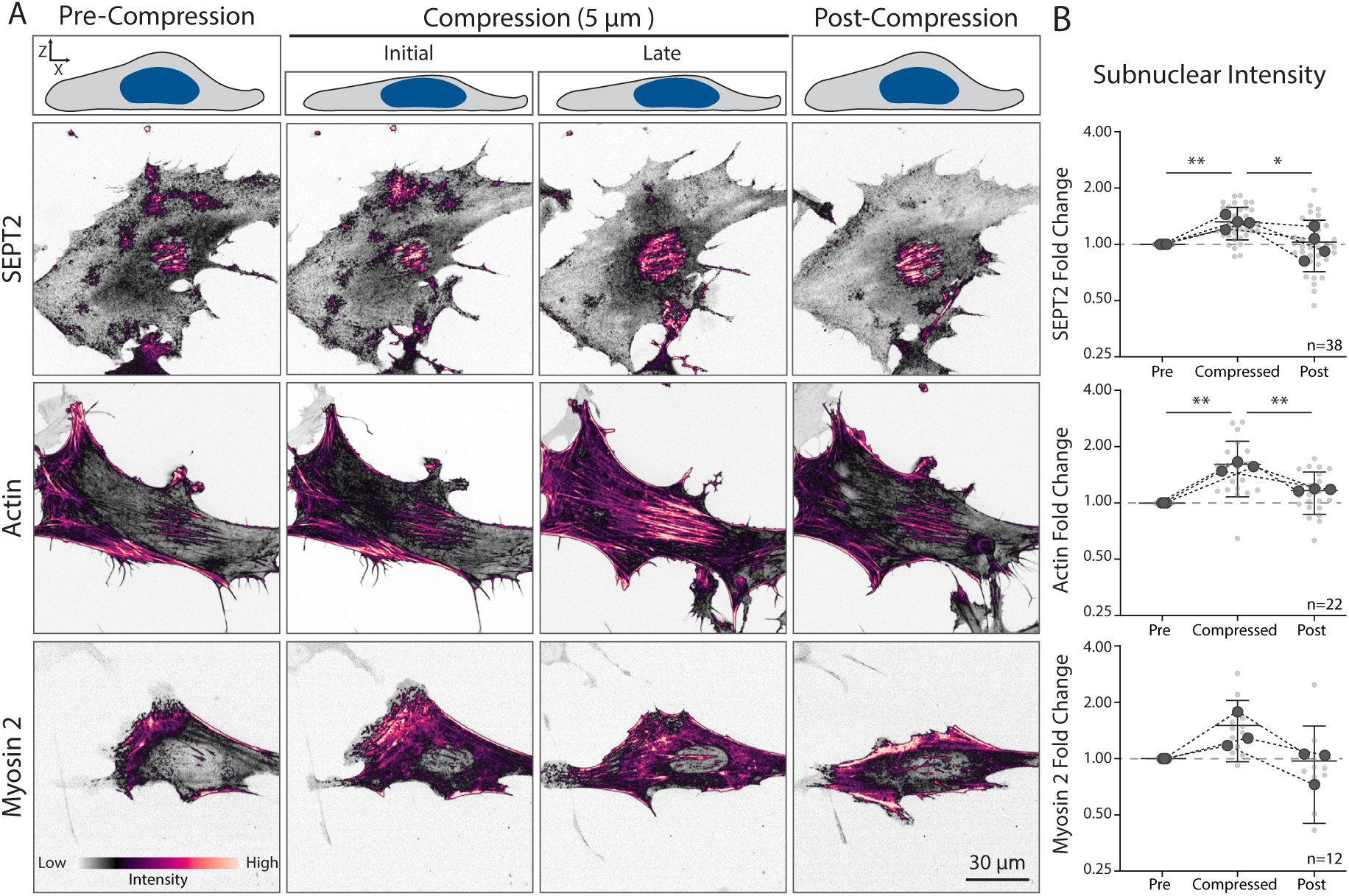
Compression increases subnuclear septin network assembly. **A)** Representative confocal images of the ventral surface of SEPT2-Halo cells with Halo dye, or expressing EGFP-FTractin or EGFP-NM2A, before, during, and after compression. **B)** The fold intensity change for septin, actin, and myosin in the subnuclear regions of the cell. Larger dots represent replicate means and smaller dots represent single cells. Bars indicate mean ± SD (n = 38 for septin, n = 22 for actin, n = 12 for NM2A). A nested 1 way ANOVA was used to determine significance with * = *p* < 0.05, ** = *p* < 0.01.

### Internally applied forces induce septin network formation around the nucleus

While our data indicates that extracellular forces can induce septin network formation, we sought to test whether internally-generated forces could elicit a similar response. We imaged SEPT2-Halo cells migrating through a microfluidic device with different features, including narrow gaps that would induce nuclear compression as the cell squeezed through [28]. We observe robust septin accumulation along the membrane as the nucleus is squeezed through points of constriction (Fig. 4A,B; Movies M3,M4). Interestingly, this behavior is not restricted to these narrow passages, as we observe septin enrichment whenever the nucleus is pushed against the walls of any of the features in the device (Fig. 4C-E; Movie M4). To quantify this, we measured the average fold increase in septin intensity along the wall as a function of the cortex distance from the nucleus. We found an average septin intensity increase of ∼ 30% when the nucleus was pushed against any wall in the device. While previous studies have shown that septins preferentially recognize and assemble on curved membranes [29–31], we found no difference in accumulation based on the geometry of the wall (Fig. S4). Together these data indicate that intracellular force production is sufficient to induce septin assembly independent of curvature.

**Figure 4.**
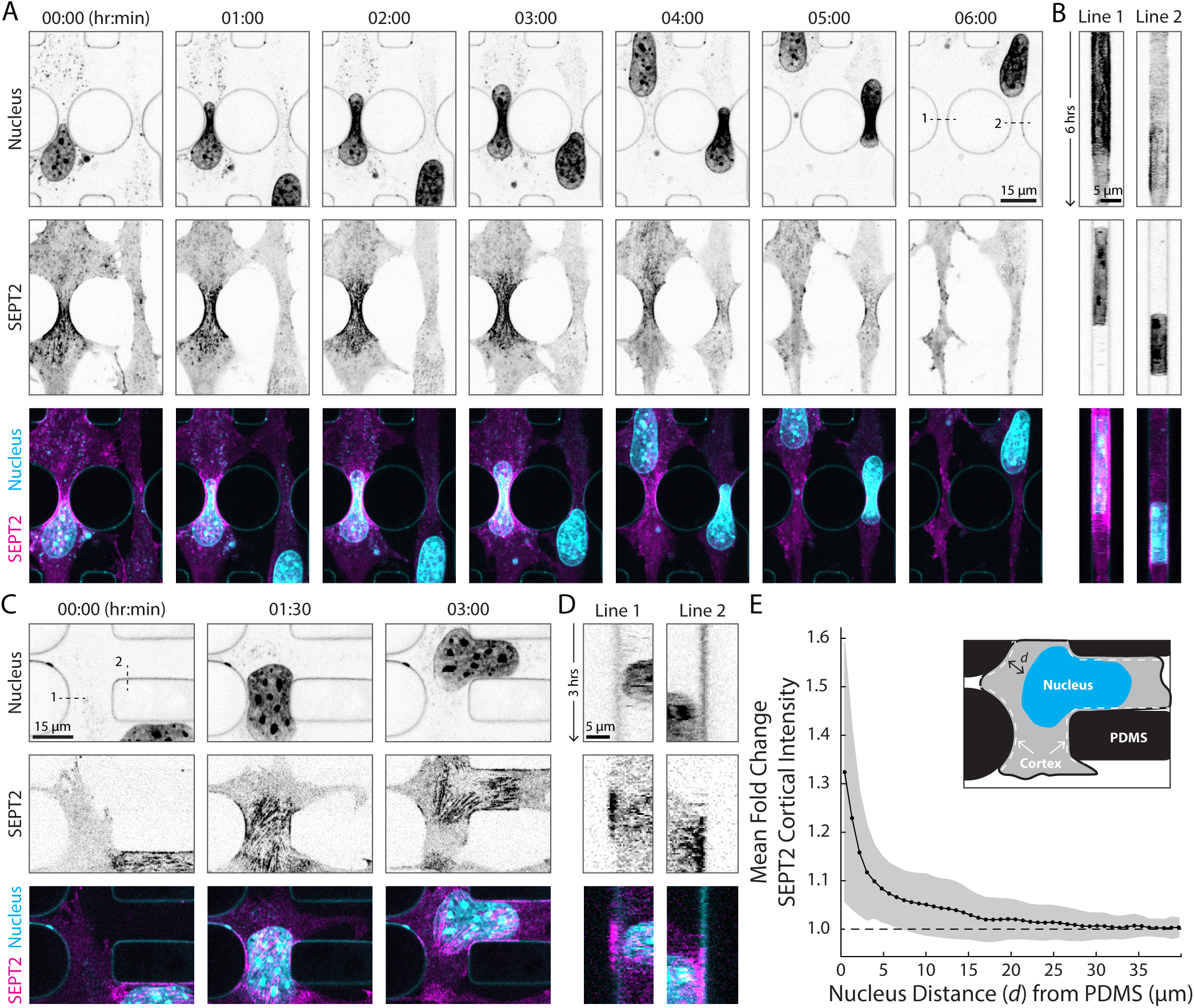
Internally generated forces induce septin assembly at cortex when the nucleus is near. **A)** Images from a timelapse (Movie M3) of SEPT2-Halo cells migrating through a microfluidic device showing septin accumulation as the cell squeezes the nucleus through narrow openings. **B)** Kymographs of the dotted lines indicated in (A) showing septin accumulation around the nucleus during migration through the constriction point. **C)** Images from a timelapse (Movie M4) showing septin accumulation occurs anywhere the nucleus pushes against the cortex. **D)** Kymographs from dotted lines indicated in (C) showing septin accumulation coincides with the nucleus nearing the PDMS features. **E)** Mean fold change in septin cortical assembly along the cortex as a function of distance from the nucleus. All data is normalized to the average SEPT2 intensity at the 30 *µ*m distance (dashed line), representing the average cortex intensity in the absence of the nucleus. Points represent the mean value and the shaded regions are the standard deviation of all points along the cortex taken from 31 different cells, across 19 fields of view, from 3 different experiments. Time in hr:min.

### Mechanical signaling drives septin network formation

We next sought to test whether the nucleus is specifically required for eliciting this septin accumulation. Cells were centrifuged in a density gradient to remove the nucleus [32], and the resulting cytoplasts were plated and imaged. Cytoplasts exhibit dynamic actin polymerization, including being able to migrate, but fail to form central septin and actin networks like their cell counterparts (Fig. 5A). To quantify this difference, we measured the alignment of septins into filaments using a previously established approach [33] and find that cytoplasts contained significantly less organized structures in the center of the cell where the perinuclear filaments typically form. To test whether the loss of these septin networks is dependent on signaling molecules associated with the nucleus, we added large glass beads with a diameter between 8-12 *µ*m into both cells and cytoplasts. Internalized beads often accumulate in the cell center near the nucleus, and the septin network assembles under both the nucleus and beads (Fig. S3). In cytoplasts we find that large glass beads at least partially rescue central septin network formation (Fig. 5C). Together these results indicate that the perinuclear septin network formation is not entirely dependent on nuclear signals and can be at least partially induced via a similarly sized large object. These findings demonstrate a clear role for mechanical signaling in mediating the formation of ventral septin networks.

**Figure 5.**
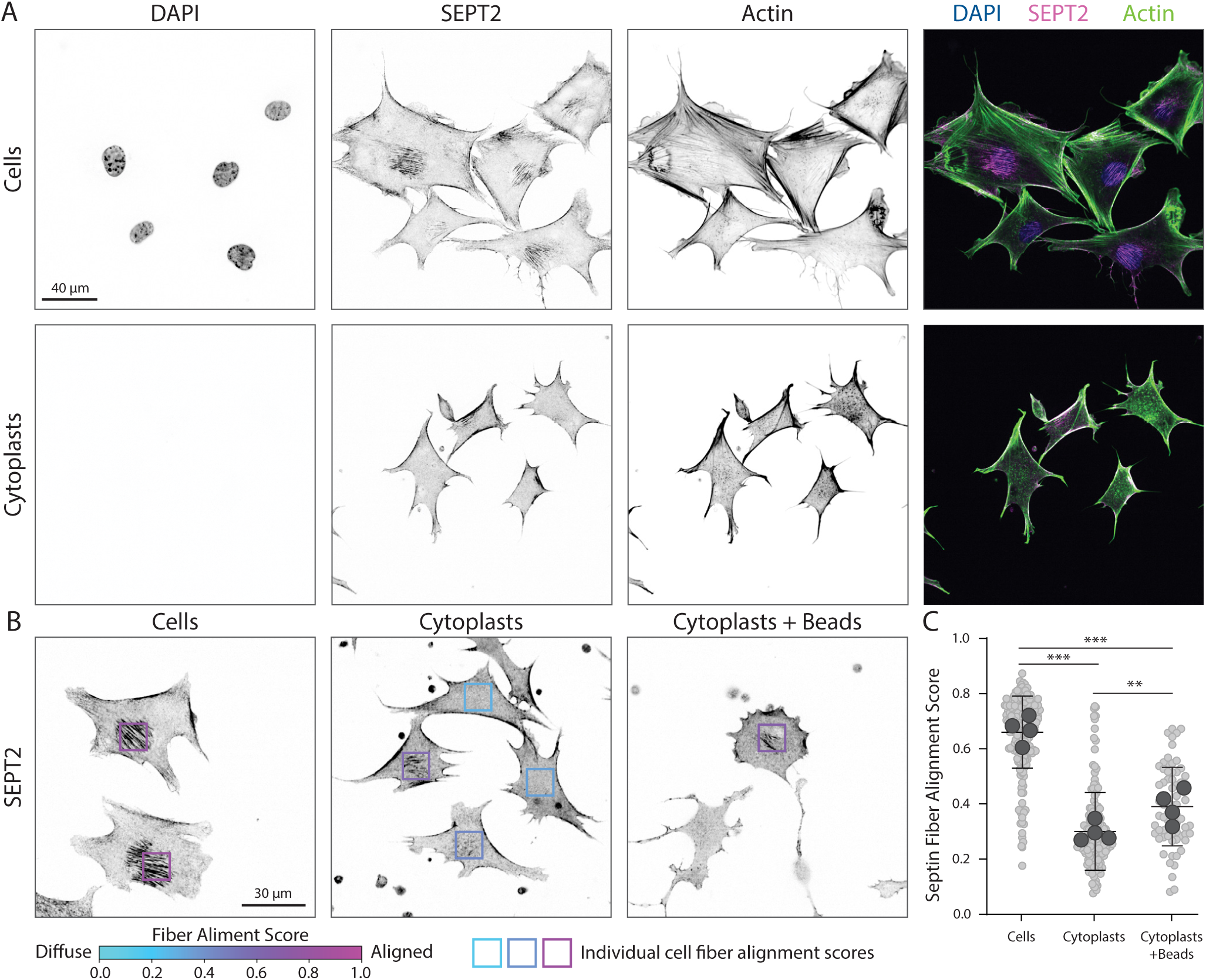
Mechanical signaling mediates ventral septin network formation. **A)** SEPT2-Halo cells and cytoplasts were fixed 2 hrs after plating and stained for actin (phalloidin) and nuclei (DAPI). **B)** Fixed images of SEPT2-Halo cells, cytoplasts or cytoplasts containing 8-12 *µ*m glass beads. Boxes indicate the regions analyzed for septin fiber formation. The box color indicates the magnitude of alignment from diffuse (0) to aligned (1) according to the range indicator below. **C)** Quantification of alignment of SEPT2-Halo cells, cytoplasts or cytoplasts with beads. Bars indicate mean ± SD (cells n = 214 , cytoplasts n = 155 , cytoplasts + beads n = 69). A one-way ANOVA was used to determine significance with ** = *p* < 0.01, *** = *p* < 0.001.

### Septins buffer the nucleus during mechanical strain

As our data suggest that septin network formation is responsive to mechanical stimuli, we sought to test the contribution of septins during confined migration. We therefore knocked down SEPT7 (SEPT7-KD) in our SEPT2-Halo cells, as SEPT7 is critical to septin octamer and hexamer formation [20]. Relative to non-targeting (NT) controls, SEPT7-KD cells have decreased expression of both SEPT7 and SEPT2 (Fig. 6A,B). Imaging of SEPT2-Halo also reveals minimal remnant septin signal and no apparent actin-septin ventral network, suggesting broad septin network disruption. We plated SEPT2-Halo and SEPT7-KD cells on transwells with 3 *µ*m pores, and compared nuclear morphology between groups before and after migration through the pores. Nuclear morphology was indistinguishable in control cells on either side of the transwell (Fig. 6C,D). In contrast, nuclei of SEPT7-KD cells that migrate through the pores exhibit deformed morphologies and significantly increased aspect ratios (Fig. 6C,D).

**Figure 6.**
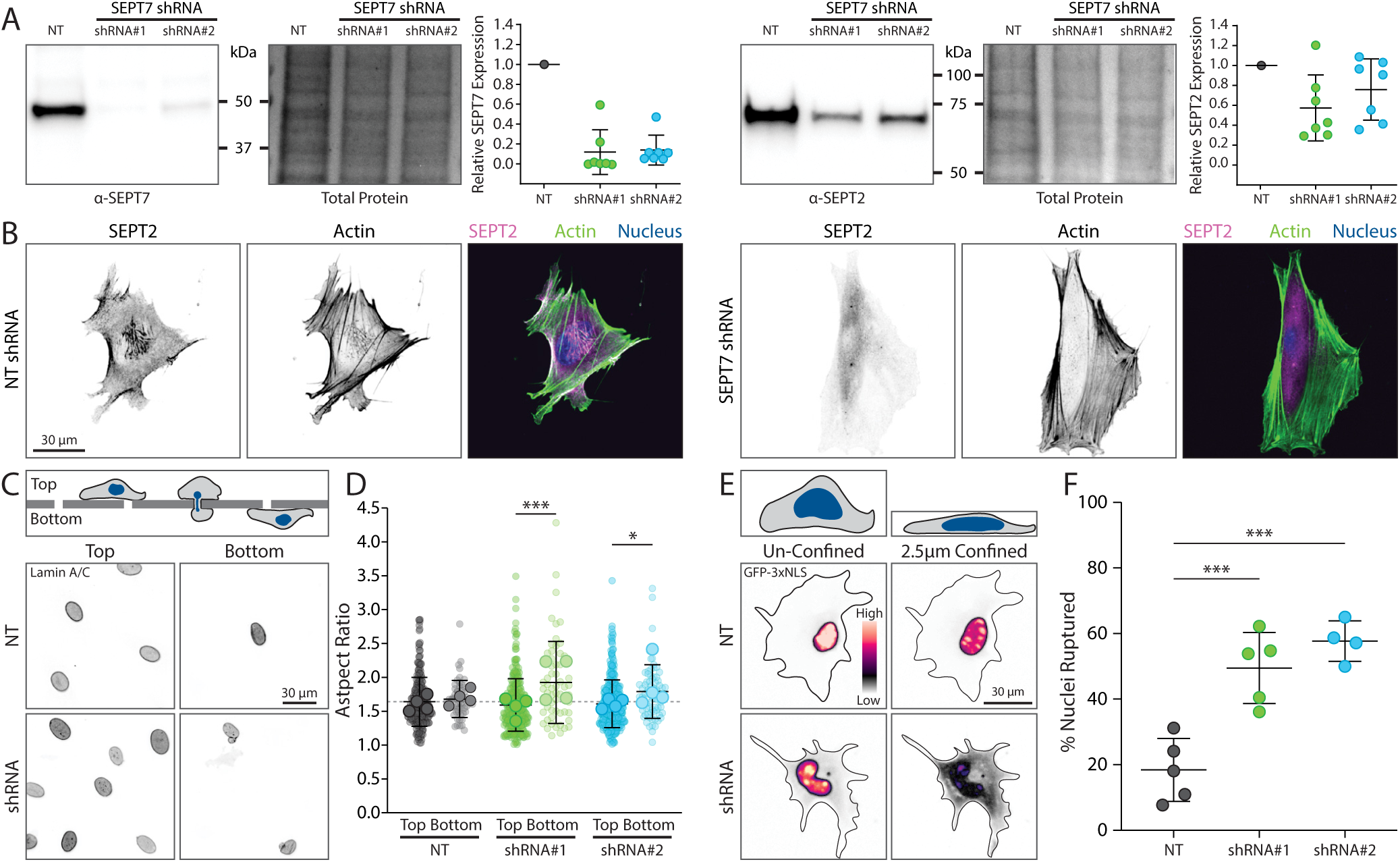
Septins buffer the nucleus during mechanical strain. **A)** SEPT2-Halo cells were treated with non-targeting (NT) or SEPT7 shRNAs. Whole-cell lysates were probed with SEPT7 (left) or SEPT2 (right) antibodies via Western blot. **B)** Fixed images of SEPT2-Halo NT (left) and SEPT7 (right) shRNA cells labeled with Halo dye and stained for actin (phalloidin). **C)** NT and SEPT7 shRNA cells were plated on a transwell with 3 *µm* pores and allowed to migrate overnight. Transwells were extracted, fixed, and immunostained for lamin A/C. **D)** Nuclear aspect ratio on top and bottom of the transwell was measured. Experimental replicates and total cells imaged for NT were 4 and top = 250, bottom = 54, for shRNA#1 were 4 and top =278, bottom= 59, and for shRNA#2 were 4 and top = 329, bottom = 77. A 2-way ANOVA was used to determine significance between top and bottom for each condition. **E)** NT and SEPT7 shRNA cells exogenously expressing 3xNLS-GFP were imaged unconfined and confined to 2.5*µ*m. **F)** The percentage of nuclei that ruptured upon confinement was measured. Circles indicate replicate means and bars indicate mean ± SD. Experimental replicates and total cells imaged for NT were 5 and 474, for shRNA#1 were 5 and 310, and for shRNA#2 were 4 and 189. A one-way ANOVA was used to determine significance. For all statistics * = *p* < 0.05, and *** = *p* < 0.001.

While we detected differences in the transwell assay, we suspected that many cells traversed overlapping pores larger than 3 *µ*m due to the random distribution of pores (Fig. S5). We therefore sought to test the response of the SEPT7-KD cells to direct compression. We exogenously expressed a 3xNLS-GFP to visualize the integrity of the nuclear envelope and measured the rate of nuclear rupture when the cells are compressed to a height of 2.5 *µ*m. While only ∼20% of control nuclei rupture at this level of compression, ∼50% of SEPT7-KD nuclei rupture (Fig. 6E,F). These data thus reveal a critical role for septins in buffering mechanical forces on the nucleus.

## Discussion

The septin cytoskeleton has been implicated in mechanotransduction, but specific contributions remain poorly defined. Here, we used endogenously-labeled SEPT2 fibroblasts to investigate the role of septins in responding to applied forces. Septins associate with a subpopulation of the actin cytoskeleton, with specific enrichment in perinuclear stress fibers (Fig. 1). We find that this septin accumulation is the result of the nucleus pushing on the cortex (Fig. 5), and that this response can result from both internally or externally generated forces (Figs. 3 & 4). Strikingly, this response can be partially replicated simply by swapping the nucleus for a glass bead, indicating a clear role for mechanics in mediating this signaling (Fig. 5). Septin knockdown subsequently perturbs nuclear morphology following migration through constricted spaces, and increases nuclear rupture upon compression (Fig. 6). Together, these data suggest that local septin accumulation reinforces the actin cytoskeleton in these regions to buffer forces applied to the nucleus.

This mechanism is consistent with previous reports showing that knockdown of septins reduces global cortical stiffness [34–37], and that loss of SEPT9_i1 in particular resulted in nuclear deformation [38]. Our data, however, suggests a dynamic spatial component to this response, helping to target septins to specific regions of the cortex that are tensed. A local reinforcement role could also explain why septins localize to other structures in the cell, such as the base of cilia [39], the sperm annulus [24], neuronal spines [40], and even the contractile ring - all regions where the cortex is likely under increased local stress. Similarly, it could explain septin accumulation at the base of blebs, where septins are thought to help stabilize the ruptured cortex as it rebuilds [41–44].

Septin’s role in buffering mechanical forces is most immediately reminiscent of a similar role ascribed to intermediate filaments [45]. Keratin, an intermediate filament, helps to maintain the mechanical integrity of epithelial sheets by anchoring the cell to the extracellular matrix through desmosomes, and disruption of this network results in blistering diseases [46]. Similarly, vimentin is an intermediate filament expressed in mesenchymal cells that contributes to cell stiffness and buffers mechanical forces to protect the nucleus [47]. Like intermediate filaments, septins are non-polar filaments that build larger networks and architectures through lateral association, but they also exhibit key differences. While intermediate filaments are distributed throughout the cytosol, with some enrichment around the nucleus [48], septins are primarily cortical. This localization, orchestrated by localized signaling (e.g. Rho family GTPases [17, 49, 50]) and binding partners (actin, membrane, phosphoinositides, etc. [29, 51–54]), could create heterogeneous subcellular mechanical support by septins, as opposed to broader cell-scale mechanical support provided by intermediate filaments. As our data further demonstrates that both extracellular forces upon the cell and intracellular forces generated within the cell can initiate cortical septin assembly, it suggests this mechanisms can act at multiple force scales. In some ways, this multimodal force response is both reminiscent of and implicative of mechanosensitive ion channels, like PIEZO, that respond to a range of mechanical forces [55]. Thus, while both intermediate filaments and septins may be mechanical buffers, we theorize the two networks are more likely parallel than redundant systems with unique responses to the spatial distribution, magnitude, direction, and kinetics of the applied force.

Exactly how septins enable nuclear mechanical resilience remains to be determined, but we speculate on some possible mechanisms and additional contributions. A simple “bed of nails” model is that perinuclear actomyosin and septin networks distribute the load more uniformly and broadly across the nucleus. Disruption of these networks results in asymmetric forces upon the nucleus, like squeezing a balloon at a single cross-section point, and leads to nuclear membrane rupture. Alternatively, our iPALM data (Fig. 2) places septins between the actin fibers and the membrane, consistent with previous data and known interaction partners [18, 29, 53, 54]. Therefore, septins could be acting as a type of lubricant during nuclear translocation, limiting nuclear engagement with cortical and plasma membrane components while allowing the actin fibers to couple to the nucleus and LINC complexes [56, 57], or potentially by altering local contractility to prevent a disrupted actin network from being shredded by myosin motors. Further, septins could be scaffolding, corralling, or sequestering cortical and transmembrane components for local signaling and structure [39, 58]. Importantly, these septin functional models are not exclusive to the nucleus, and could be happening with smaller organelles impinging upon the cortex and plasma membrane that are less apparent due to the magnitude of response. More than likely, multiple of these potential mechanisms are occurring in parallel.

Functionally, septins are strongly implicated in regulating immune cell migration [34, 59]. As immune cells must navigate dense tissue and extracellular matrix as they traverse the body, their nucleui are constantly being squeezed, and thus enrichment of septins in the cortex of these cells is consistent with our findings here. It is thus also similarly unsurprising that up-regulation of septin expression is associated with a number of cancers [60–63]. In work submitted concurrently with this manuscript, Mayca-Pozo and colleagues flesh this idea out explicitly [64], showing that septins protect against large-scale genomic alterations (or excessive genomic instability) during confined migration, and that over-expression of septins could be mechanoprotective for cancer cells that would otherwise succumb to apoptotic signals. These results, combined with our data, suggest that increases in septin expression could therefore facilitate metastasis by specifically enabling cells to migrate resiliently through confined spaces. Interestingly, methylation of septin is a proposed biomarker for tumor detection and treatment progress [65–67], suggesting that post-translational modifications, and not just expression, could contribute to this mechanical response mechanism.

In conclusion, our data reveal a novel role for septins in mediating mechanical signaling in the cell, specifically around the nucleus. This behavior is reminiscent of, but distinct from, the role of intermediate filaments, and appears focused on dynamically and locally reinforcing the cell cortex. This mechanism acts to protect the nucleus from damage, and facilitates cell migration through confined spaces. Coupled with the work of Mayca-Pozo et al. [64], our results shine a spotlight on a critical role for septins in mechanotransduction, where they coordinate mechanical signaling in the cortex, and prevent damage to the nucleus. There is still much to learn about the various functions of septins as they continue to demonstrate that their role extends well beyond their function in cell division.

## Methods

### Cell culture and Transfections

Immortalized mouse embryonic fibroblasts (MEF) were a kind gift of Dr. Mary Beckerle’s lab (University of Utah, Salt Lake City, UT). HEK293FT cells (CRL-3216; ATCC) were cultured for making lentivirus. All cell lines were cultured in Dulbecco’s modified Eagle’s Medium (MT10013CV; Corning) supplemented with 10% FBS (10437-028; Gibco), 1% Antibiotic-antimycotic (#15240062; Thermofisher), and 5 *µ*g/mL prophylactic Plasmocin (ant-mpp; InvivoGen). All cells were grown in uncoated plastic tissue culture dishes (229661, 229621; CELLTREAT). For imaging cells were plated on #1.5 coverslips (Corning) or glass bottom dishes (Cellvis) coated with 10 *µ*g/mL fibronectin (FC010; EMD Millipore) for 1 hr at 37°C. For EGFP-Ftractin [68], EGFP-NM2A (Plasmid #11347; Addgene), 3x-NLS-GFP (Plasmid #58468; Addgene) and mEmerald-LaminA (Plasmid #54139; Addgene) exogenous transfections were performed using Neon electroporation (Invitrogen) with 1x 20 ms pulses of 1600V and 3.5 *µ*g DNA per 200,000 cells in 100 *µ*L of reaction.

For lentiviral production, HEK293FT cell were plated in 6 cm dishes at 60-80% confluent. Transfection complex was generated using LipoD293 (SL100668; Signagen), packaging plasmid psPax (Plasmid #12260; Addgene), envelope plasmid pmD2.G (Plasmid #12259; Addgene), and lentiviral constructs (shRNAs TRCN0000101845, TRCN0000101847, SHC016; Sigma Aldrich). The complex was added dropwise to the HEK cells and allowed to incubate 24 hrs. Media was changed and discarded at 24 hrs, and collected at 48 and 72 hrs. Viral media was collected and filtered using 0.45 *µ*m Basix syringe filter (13-100-107; Thermofisher) SEPT2-Halo cells were infected with viral media. Viral media was removed after 24 hrs. Cells were selected with 2 *µ*g/mL Puromycin (61-385-RA; Corning).

### Generation of CRISPR knock in cell line

SEPT2-Halo knock-in cells were derived from the immortalized MEF line using CRISPR/Cas9 as described previously [50]. Briefly, we generated pSpCas9(BB)-2A-Puro (PX459) V2.0 (Plasmid #62988; Addgene) with target sequence 5’-AAACTTCA-TCAATAACCCGC-3’ using established protocols [69]. To generate donor plasmids, pUC57 was digested with EcoR1 and Stu1 and purified. A four-piece Gibson assembly was then performed with three gBlocks (IDT): (1) a 794 bp 5’HDR of genomic sequence immediate upstream of the endogenous start codon, (2) HaloTag with an 18 amino acid GS-rich linker, (3) an 802 bp 3’HDR of genomic sequence immediately downstream of the endogenous start codon with silent PAM site mutation. Fibroblasts were transfected with donor and target-Cas9 plasmids and single-cell sorted. Individual clones were evaluated for knock-in via Western blotting and microscopy.

### Microscopy

Unless stated otherwise (e.g. SIM, iPALM) all imaging was performed with a Plan-Apochromat 63X/1.4 NA objective or a Plan-Apochromat 63X/1.46 NA TIRF objective on Axio Observor 7 microscopes (Zeiss) equipped with a CSU-W1 Spinning Disk (Yokogawa), Prime 95B sCMOS cameras (Photometrics), and enclosed in a temperature and humidity regulated environmental chamber (Oko Labs) controlled by Slidebook Software (Intelligent Imaging Innovations). For excitation of fluorophores, 405, 488, 561, and 637 nm lasers were used.

### Fixation

Cells were fixed using a 4% paraformaldehyde solution (15710; Electron Microscopy Sciences) in cytoskeletal buffer (CB) (0.1M MES, 0.03M MgCl_2_, 1.38M KCl, 0.02M EGTA in 1L H_2_O) for 15 min at 37°C. The PFA solution was removed, cells were rinsed with 1xPBS, permeabilized with a solution of 0.5% Triton X-100 (Fisher BioReagents) in CB for 15 min, rinsed in 1xPBS, and blocked with a solution of 1.5% bovine serum albumin (BSA) (CAS 9048-46-8; Research Products International) in CB. Fixed samples were incubated with primary antibodies at 4°C overnight, washed in 0.05% Triton X-100 in 1xPBS for 3×10 min with gentle rocking, incubated in secondary antibodies diluted in blocking solution for 1 hr at room temperature, and washed in 0.05% triton in PBS 3×10 min before imaging or mounting. Cells were mounted using ProLong Glass antifade mountant (P36980; ThermoFisher). Mounted samples were allowed to cure 18 hrs in a dark drawer at room temperature prior to imaging.

### Antibodies and Dyes

The following antibodies and dyes were used for western blotting (WB) and immunofluorescence (IF) and live imaging: Anti Lamin A/C (39087; Active Motif; IF 1:500). Anti SEPT2 (11397-1-AP; Protein Tech; IF and iPALM 1:500). Anti SEPT2 (60075-1-Ig; Protein Tech; WB 1:2000). Anti SEPT7 (13818-1-AP; Protein Tech; WB 1:2000). Anti SEPT7 (18991; IBL America; IF 1:500). Anti SEPT9 (A8657; ABclonal; IF 1:500 and WB 1:2000). Goat Anti Rabbit AlexaFluor 488 (A32731; ThermoFisher Scientific; IF 1:1000). Goat Anti Mouse AlexaFluor 488 (A32723; ThermoFisher Scientific; IF 1:1000). Goat Anti Rabbit AlexaFluor 647 (A21244; ThermoFisher Scientific; IF 1:1000). Phalloidin, CF(R)583R (BOT-00064-T; Biotium; iPALM 1:200. Phalloidin-Atto 647N (65906; Sigma-Aldrich; IF 1:5000). DAPI (D1306; ThermoFisher Scientific; IF 1:5000). Hoechst (H3570; Invitrogen; live imaging 1:5000 for 10 min). SPY650-FastAct_X (CY-SC502; Cytoskeleton; live imaging 1:1000). Janelia Fluor JFX554 HaloTag Ligand (HT103A;Promega; live imaging 10-50nM) Janelia Fluor 646 HaloTag Ligand (HT106A;Promega; live imaging 10-50nM)

### Collagen gels & Polyacrylamide hydrogels

Collagen gels were generated using a protocol adapted from [70]. 3.0 mg/mL rat tail collagen (354236; Corning) was mixed with cell culture DMEM and reconstitution buffer on ice. 200-250 *µ*L of the collagen mixture was added to pre-chilled 35 mm glass bottom dishes (Cellvis). Dishes were pre-coated with Poly-D-Lysine (A3890401; Gibco) for 10 min at room temperature. Gels were allowed to polymerize at 18°C overnight. Cells were plated on top of gels in media with Halo dye and allowed to invade for 24 hrs before imaging. Halo dye was removed 1 hr before imaging.

Polyacrylamide gels were made as previously described [71]. Briefly, gels were polymerized on an activated coverslip with 7.5% acrylamide and 0.3% bis-acrylamide for a shear modulus of 8 kPa. The gels were covalently coupled with fibronectin (1 mg/mL) via sulfo-sanpah (NC2028693; Fisher Scientific).

### Septin localization along nucleus and cell centroid axis

Wild-type MEFs were fixed and immunostained for SEPT2 and DAPI-stained for nuclei. The SEPT2 channel was used to generate a cell mask and identify the cell centroid. The nuclear channel was used to generate a nuclear mask and identify a nuclear centroid. To subtract background in the SEPT2 channel, the intensity of a large region outside of the cell was measured and subtracted from the entire image. A bounding box was manually traced around the prominent septin medio-ventral network and the center of mass was measured in the background-subtracted SEPT2 channel. The distance of the septin center of mass along the nuclear cytoplasmic axis was calculated and presented as fraction of the distance between the cell centroid (value of 0) and the nuclear centroid (value of 1), where positive values indicate a septin preference towards the nucleus and negative numbers indicate a septin preference away from the nucleus.

### Structured Illumination Microscopy

SEPT2-Halo cells were transiently transfected with mEmerald-LaminA and stained with Spy650-FastActin_X (CY-CS502; Cytoskeleton). SIM imaging was performed on a Zeiss Lattice SIM 5 with a Plan-Apochromat 63x/1.40 NA oil objective and Hamamatsu Fusion BT cameras. Z-stacks were collected in Leap mode with optimal step size. Processing was performed in ZEN BLUE 3.9 with SIM^2^ and the following parameters: result sampling = 2; processing sampling = 2; auto sharpness = on; order combination = Wiener filter; regularization = none; iterations = 3 for actin and septin, 2 for laminA; fitting = fast fit; histogram = scaled to raw.

### iPALM

iPALM imaging and analysis was performed similar to previous work [72–76]. For sample preparation, MEFs were plated on 25 mm diameter round coverslips containing gold nanorod fiducial markers (A12-40-600-CTAB; Nanopartz, Inc.), passivated with a ∼50 nm layer of SiO_2_, deposited using a Denton Explorer vacuum evaporator, except the coverslips were not coated with any ECM protein before plating the cells. Coverslips were incubated in complete cell culture media for 1 hr at 37°C prior to plating cells. Cells were plated for ∼18 hr before fixation as above. Rinse steps are different than listed above, 6 sequential washes in PBS were performed (3×5 min, 2×10 min). Actin was labeled using phalloidin-CF583R (#00064-T; Biotium). Septins were labeled with either anti-SEPT2 or a combination of anti-SEPT2 and anti-SEPT7 to achieve higher labeling density, followed by goat-anti-rabbit AlexaFluor647 secondary antibody. The coverslips were then covered with STORM buffer, prepared as follows. 17 mg/mL Catalase (C100-50MG, Sigma-Aldrich) and 70 mg/mL Glucose Oxidase (G-2133-50KU, Sigma-Aldrich) solutions were prepared in Buffer A (10 mM Tris pH 8.0 + 50 mM NaCl). Both were mixed in a 1:4 volume ratio to prepare GLOX solution. This was centrifuged and only the supernatant was used. 1 M cysteamine (30070-50G, Sigma-Aldrich) solution was prepared in 0.25 N HCl. The STORM buffer was prepared by combining GLOX solution, 1 M cysteamine, and Buffer B (50 mM Tris pH 8.0 + 10 mM NaCl + 10% Glucose) in 1:10:90 volume ratio. Following coverslips being covered with STORM buffer, an 18 mm diameter round coverslip (CS-18R17, #1.5 thickness; Warner Instruments) was placed centrally on top of the sample and the boundary was sealed with Valap (1:1:1 of vaseline, lanolin, and, paraffin) to create a sandwich mounting. The mounted samples were then imaged on the iPALM.

For imaging, actin samples were excited using 561 nm laser (Opto Engine LLC) excitation at 1–2 kW/cm2 intensity in TIRF conditions. 100,000 images were collected through dual Nikon Apo TIRF 60x/1.49NA objective lenses coupled to a 593/40 nm bandpass filter (Semrock), and acquired via three EMCCD cameras (DU987U; Andor) with 50 ms exposure. Images were XYZ drift and tilt-corrected, channel-merged, and checked for alignment quality using the PeakSelector software (Janelia Research Campus80). Alexa Fluor 647-labeled septin was imaged similarly, but with 2–3 kW/cm2 intensity 647 nm laser excitation (Opto Engine, LLC.), 647 nm long-pass filter (Semrock), and 30–40 ms exposure time.

For analysis, the iPALM data was processed/localized using PeakSelector software (Janelia Research Campus). The gold fiducials embedded in the coverslip were used for drift correction. iPALM data were analysed using iPALM plotter (AIC, Janelia Research Campus; https://github.com/aicjanelia/ipalmplotter) and images were rendered using the PeakSelector software. iPALM localization data records both the fluorescent molecules localized within the fibers, as well as molecules in the cytoplasmic fraction. iPALM images were rendered using PeakSelector. To quantify the spatial distribution of the proteins within individual fibers, we identified fibers using custom MATLAB code (https://github.com/aicjanelia/iPALM-fiberAnalysis). After the gold nanorod fiducial markers were removed from the rendered images using custom MATLAB code (https://github.com/aicjanelia/iPALM-beadRemoval), localizations were roughly pruned by excluding localizations at unreasonably large z-positions—likely ghosting artifacts. A rendered image was produced by counting localizations within pixels of a given size. Fibers were detected in this rendered image through a series of morphological processing steps: MATLAB’s fibermetric filter, binary opening and eroding, and skeletonization. Connected component analysis of the resulting binary image detected fibers. Small fiber segments were discarded at this step. Fibers were then iteratively segmented into short linear pieces. Localizations within a constant width of these segments were assigned to the fiber. By iterating over all segments for each fiber, complete sets of fiber-associated localizations were determined. The z-position of these localizations was subsequently recorded along the length of the fiber. This process was repeated for each channel. Fibers with low or non-uniform density and fibers not parallel with the coverslip were removed. For analysis of protein distributions in fibers, the three-dimensional molecular coordinates for each region (individual fibers) were analyzed to extract the median axial height for actin and septin, with the delta of actin minus septin plotted accordingly.

#### Western Blotting

Cells were collected, counted and pelleted. Complete Laemmli buffer was made using 2x Tris-Glycine Laemelli sample buffer (1610737; Biorad) with 10% beta-mercaptoethanol and protease and phosphatase inhibitor cocktail (78442; ThermoScientific). 100 *µ*L of complete buffer was added per 500,000 cells in a centrifuged pellet. Samples were boiled for 5 minutes at 95°C, and cell lysates were loaded into 4-15% Mini-PROTEAN TGX Stain-Free Gels (4568085; Bio-Rad). Protein was transfered using a Trans-Blot Turbo system (Biorad) onto a polyvinylidene fluoride (PVDF) membrane. Blots were stained with Amido Black (CAS 1064-48-8; Bio Basic) for 10 min and rinsed in de-stain buffer (5% acetic acid in water) for 3×5 min. Total protein from blots was imaged on a chemidoc (Biorad). Blots were blocked for 1 hour at room temperature (RT) in 3% BSA in PBS with 0.1% Tween (PBS-T). Primary antibodies were added in blocking buffer and rocked at 4°C overnight. Blots were washed in 0.1% PBS-T 3×10 min, incubated in secondary antibodies for one hour at RT, washed in 0.1% PBS-T for 3×10 min, incubated incubated in Clarity Western ECL Substrate (170561, Bio-Rad) for 1 min and imaged. Blots were analyzed using FIJI, samples were background subtracted and normalized to total protein. Normalized values were then compared to WT to determine relative expression values.

### Microfluidic devices

The microfluidic chamber design was adapted from previously used designs [77], and a corresponding SU8 wafer for making the molds was fabricated by the UIC Nanotechnology Core Facility. Briefly, a negative of the wafer was cast in PDMS (Sylgard 184 Silicone Encapsulant, 50-366-794; Fisher Scientific), then used to create an epoxy mold. The mold was cleaned with isopropanol and dried with compressed air before use. A 10:1 mixture of Sylgard 184 base and curing agent was poured in the epoxy mold, vacuum-degassed for 30 minutes, and baked at 70°C overnight. The PDMS was allowed to cool before demolding. Holes were then punched in the PDMS for the media container (2.5mm) and cell port (1mm). The features were cleaned with Scotch tape, washed twice with 70% ETOH, twice with MiliQ water, and dried with compressed air. To assemble the microfluidic device, the PDMS slab was coupled immediately to a 35 mm glass bottom dish (Cellvis) after both were plasma cleaned (Harrick Plasma) for 30 seconds. The coupled device was then heated at 70°C for one hour to strengthen the bond.

Before using, the microfluidic device was cooled to room temperature, plasma treated for 2 min, and coated with 20 *µ*g/ml fibronectin at 37°C for 20 min. It was then rinsed with PBS and filled with cell media and kept at 37°C for 2 hr. Cells were resuspended at 2.5 million/ml, and 10 *µ*l of the cell suspension was inserted into the cell port. The cells were allowed to attach for 5-10 minutes, prior to filling the media port with media. An additional 4 mL of media containing 1 *µ*g/ml Hoescht was added to flood the device and prevent evaporation. The device with cells was incubated overnight at 37°C with 5% CO_2_, then moved to the microscope and imaged every 2 minutes under the same conditions.

To quantify septin enrichment at the cortex, images were registered using the brightfield channel and the calculated translation was applied to the other channels. A binary mask of the PDMS features in the microfluidic device was made and regions of interest that defined an area over a single cell were defined manually. Objects in the feature mask were then outlined to create a list of pixels representing the surfaces of the objects. For each frame the nucleus image was thresholded to define the nucleus boundary, and a distance map from the binary nucleus mask was made indicating the distance of each pixel in the image from the nucleus. For each frame, the septin intensity and the distance from the closest pixel of the nucleus was recorded for the list of points defining the outlines of the PDMS objects. Points were binned by distance with a bin width of 5 pixels and then averaged over all points in the movie. Intensity values were normalized to the average intensity at 30 *µ*m, and plotted as a function of the distance of the nucleus from the PDMS. The plot in Fig. 4E represents the average of 31 cells from 19 fields of view, taken across 3 different experiments. To break the data down by the curvature of the PDMS features (Fig. S4) masks of only the flat edges of the rectangular pillars (*κ* ∼ 0), the large round pillars (*κ* ∼ 0.1 *µm*^−1^), or the corners of the rectangular pillars (*κ* ∼ 0.35 *µm*^−1^) were used to create the point list. All analysis was performed in python and is available at https://github/OakesLab/.

### Dynamic Confinement

5 and 2.5 *µ*m pillars were made from SU8 wafer masks fabricated in the UIC Nanotechnology Core Facility. Masks were cleaned with water, dried with an air gun, and rinsed in isopropanol. 5 g of PDMS (Sylgard 184 Silicone Encapsulant, 50-366-794; Fisher Scientific) and its accompanying curing agent were mixed at a ratio of 10:1 and poured over the cleaned mask. PDMS was degassed in a vacuum chamber to remove bubbles for 15 min. 10 mm glass coverslips were plasma cleaned (Harrick Plasma) for 5 min on high. Coverslips were pushed into the PDMS covered mask and cured at 98°C for 30 min. PDMS was allowed to cool to room temperature for 10 min before peeling excess PDMS from the mask. Isopropanol was sprayed on the mask to help remove coverslips. Pillars were then dried, rinsed in PBS, and placed in a dish with complete cell media overnight in the cell culture incubator. Cells were plated in a 35 mm dish with a 28 mm glass bottom of #1.5 glass (Cellvis) coated with fibronectin (10*µ*g/mL) for 1 hr at 37°C. Cells were placed in 700 *µ*L of media for imaging at 37°C and 5% *CO*_2_. The dynamic confinement cup was a kind gift of the Piel Lab (Institut-Curie). The cup was cleaned with ethanol, allowed to dry and then tape was used to remove any dust. The pillars were attached to the piston in the center of the cup, and the entire device was coupled to the vacuum (Flow EZ; Fluigent). The cup was placed on the dish and pressure was applied to the outer edge to couple the cup to the dish. Cells were imaged at steady-state vacuum (≈ -15 mBar) for 8 min, vacuum was increased (-90 mBar) to bring down the piston and cells were imaged for 30 min. The confinement was then released by reducing the vacuum and cells were imaged for an additional 30 min.

To quantify changes in subnuclear intensity of cytoskeletal elements a standard sized circular region of interest (ROI) 25 *µ*m in diameter was marked under the centroid of nucleus. The background subtracted mean intensity of the ROI was measured at each time point, and the pre-compression intensity was taken as the average of the pre-compression values. A rolling average of 8 frames (to match the length of the pre-compression data) was calculated and the maximum intensity during compression was determined. The last 8 frames of the post-compression time points were averaged to create the post-compression mean. All three means were then normalized to the pre-compression mean providing a relative fold change.

To quantify nuclear envelope integrity upon 2.5 *µ*m confinement, nuclear and cytoplasmic masks were generated using the EGFP-3xNLS signal at varying threshold levels. These masks were applied to the raw images and nuclei were considered to rupture if the cytoplasmic signal increased following compression.

### Enucleating cells

Enucleation was performed as previously described [32]. A stock Ficoll solution was formed by dissolving Ficoll-400 (26873-85-8; BioBasic) at 50% (wt/vol) in sterile PBS without Ca^2+^/Mg^2+^ overnight while being rotated at RT. The solution was filtered (0.4 *µ*m) and stored at 4°C. This stock Ficoll was used to prepare Ficoll-DMEM stocks of 30%, 20%, 18%, and 15%. Cytochalasin-D (100-0556; StemCell Technologies) was added to each stock at a final concentration of 10 *µ*M. A discontinuous iso-osmotic gradient was poured into a 13.2 mL centrifuge tube (C14277; Beckman Coulter) from the Ficoll-DMEM stocks in the following order: 2 mL of 30%, 20% and 18%, then 1 mL of 15%. The gradient was allowed to equilibrate overnight in the tissue culture incubator. SW41 Ti rotor buckets were also incubated at 37°C overnight. 2×10^7^ dye-labeled SEPT2-Halo cells were lifted, pelleted, resuspended in 1 mL of pre-warmed 15% Ficoll-DMEM and layered on top of the gradient. DMEM was added to fill the tube. The gradient was placed into the SW41 Ti rotor bucket and incubated in the cell culture incubator for 45 min. The rotor was centrifuged at 125,000 ×*g* for 1 hr at 30°C in a Beckman Coulter Optima XL-100K Ultracentrifuge and stopped with minimal breaking. Following centrifugation, a thin band containing cytoplasts could be observed in the gradient and fractions around this band were collected. Collected fractions were washed and centrifuged twice with PBS and twice in DMEM. Control cells were incubated in 15% Ficoll-DMEM stock in cell culture incubator for the duration of the centrifugation and washed the same as the cytoplast fractions. Control cells and cytoplasts were plated in 4-well glass bottom dishes and allowed 1 hr to recover and adhere to the dish. Cells were then fixed as above. For experiments using the solid glass 8.5-12 *µ*m microspheres (P2015SL; Cospheric), microspheres were sterilized in a UV oven (UVP Crosslinker CL-1000, Analytik Jena ) at max power for 5 min. Beads were further sterilized with 70% ethanol, rinsed 3x 1xPBS, coated in PolyD-Lysine for 10 min at RT with gentle agitation, and rinsed 3×5 mL of sterile 1xPBS. Beads were added on top of adhered cells or cytoplasts, centrifuged (5810R; Eppendorf) in a custom holder for 4 well dish for 20 min at 200 rpm. Media was exchanged to remove excess beads, and cells were incubated for 2 hrs prior to fixation as above. Cells were imaged using Z-stacks with steps of 0.5*µ*m to capture the entire cell or cytoplast. To quantify the filament intensity and alignment we used the Alignment by Fourier Transform method [33]. A standard square ROI (12×12 *µ*m) was placed under the centroid of the nucleus, center of the cytoplast, or under the centroid of the bead for analysis.

### Transwell

Transwell inserts (3402, Corning) were coated in 50 *µ*g/mL collagen (354236, Corning) for 1 hr at 37°C and then rinsed in 1xPBS. Media with 0.1% FBS (Fetal Bovine Serum) and Halo dye was loaded into the top and bottom of the transwell. Cells were lifted and re-suspended in the 0.1% FBS media and plated on the top of each transwell. Cells were allowed to adhere for 6 hrs before the media in the bottom of the well is swapped for media with 10% FBS and Halo dye to generate an FBS gradient. Cells were allowed to migrate through the pores overnight. Transwells were fixed the next morning following the fixation protocol above. After blocking, the transwell membrane is cut out from the device and placed in primary antibody solution overnight. The insert was rinsed, placed in secondary antibody solution, and mounted between two #1.5 22×30 mm coverslips following mounting protocol above. Samples were either imaged as a full stack all the way through the transwell membrane or each side of the membrane was imaged separately using smaller stacks. Max projections of the lamin A/C channel for the top and bottom of the well were created in FIJI and thresholded to create a mask of all the nuclei. The analyze particle feature in FIJI was used to identify nuclie and measure aspect ratio.

## Acknowledgements

We thank Elias Spiliotis (University of Virginia), Ryan Petrie (Drexel University) and members of the Beach and Oakes labs for helpful comments, the Lammerding lab (Cornell University) for assistance with the microfluidic devices, the Piel lab (Institut Curie) for assistance with the dynamic confiner, the UIC Nanotechnology Core Facility for microfabrication, and the Loyola University Chicago Core Microscopy Facility. iPALM imaging was performed in collaboration with the Advanced Imaging Center (AIC) at Janelia Research Campus, which is supported by the Howard Hughes Medical Institute. This work was supported in part by National Science Foundation CAREER Award 2000554 to P.W.O, National Institutes of Health grants R35-GM138183 to J.R.B, R01-GM148644 to P.W.O., and shared instrument grant S10OD034431 for Zeiss SIM5 to Loyola University Chicago.

## Author Contributions

MEU, JRB and PWO developed the project. MEU, AC, SC, JJT performed experiments. MEU, JJT, JRB, and PWO analyzed the data. MEU, JJT, HB, RML, OP, SK, JA, TLC, JRB and PWO performed iPALM imaging and analysis. MEU, JRB, and PWO wrote the initial manuscript. All authors edited and approved the final text.

## Supplemental Data

### Supplemental Figures

**Figure S1.**
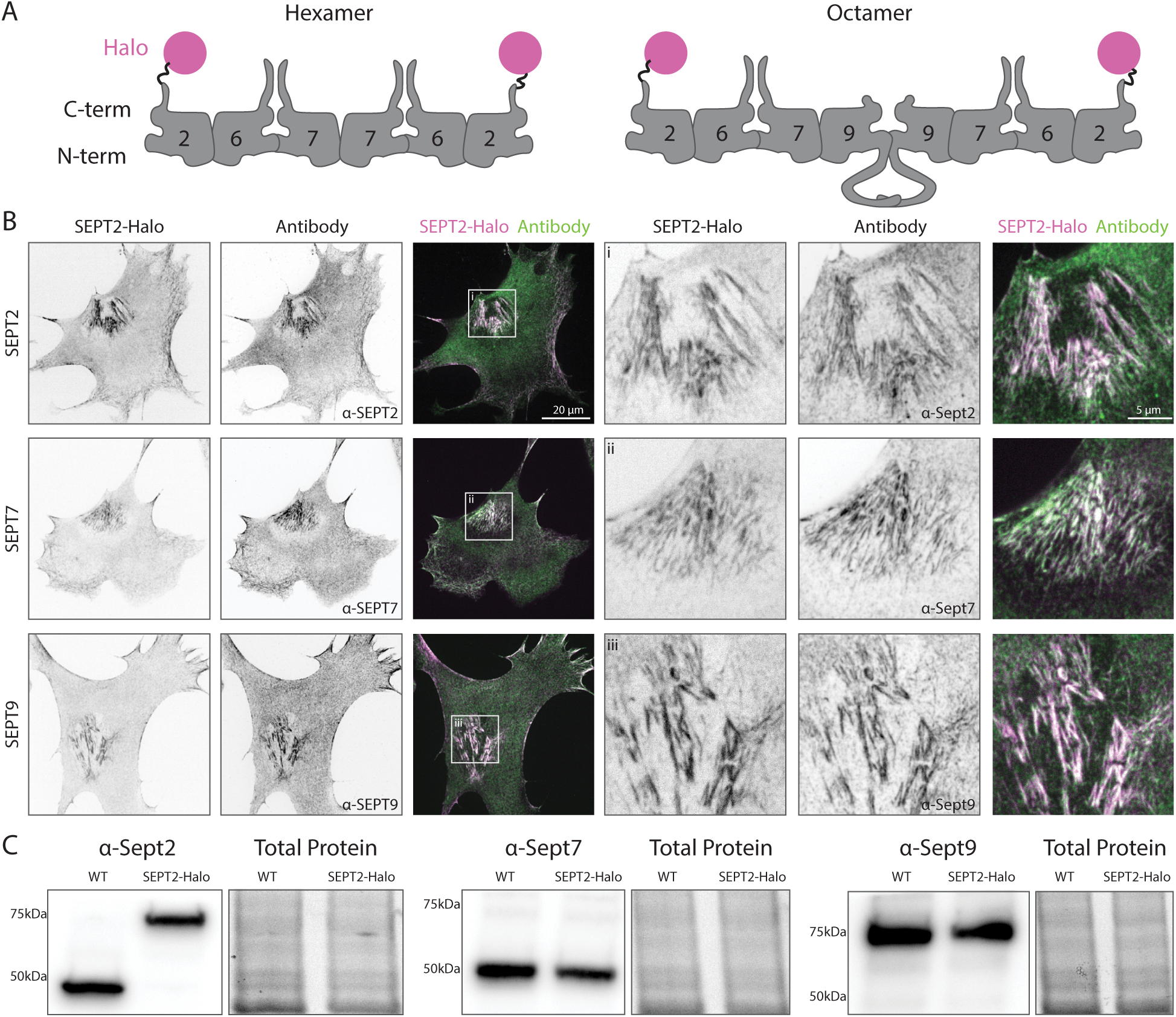
SEPT2 Knock-in cells have normal septin localization and expression. **A)** Cartoon depicting a septin hexamer and octamer with an approximation of our HaloTag and linker. **B)** Our SEPT2-Halo knock-in cells fixed and labeled with Halo dye (magenta) and immunostained for either SEPT2, SEPT7, or SEPT9 (green). Zoomed insets of the subnuclear region are shown to the right. **C)** Western blots of the parental MEF line and the SEPT2-Halo cells validating the knock-in did not change septin expression. The SEPT2-Halo band is shifted approximately 30 kDa larger, consistent with the approximate size of a HaloTag.

**Figure S2.**
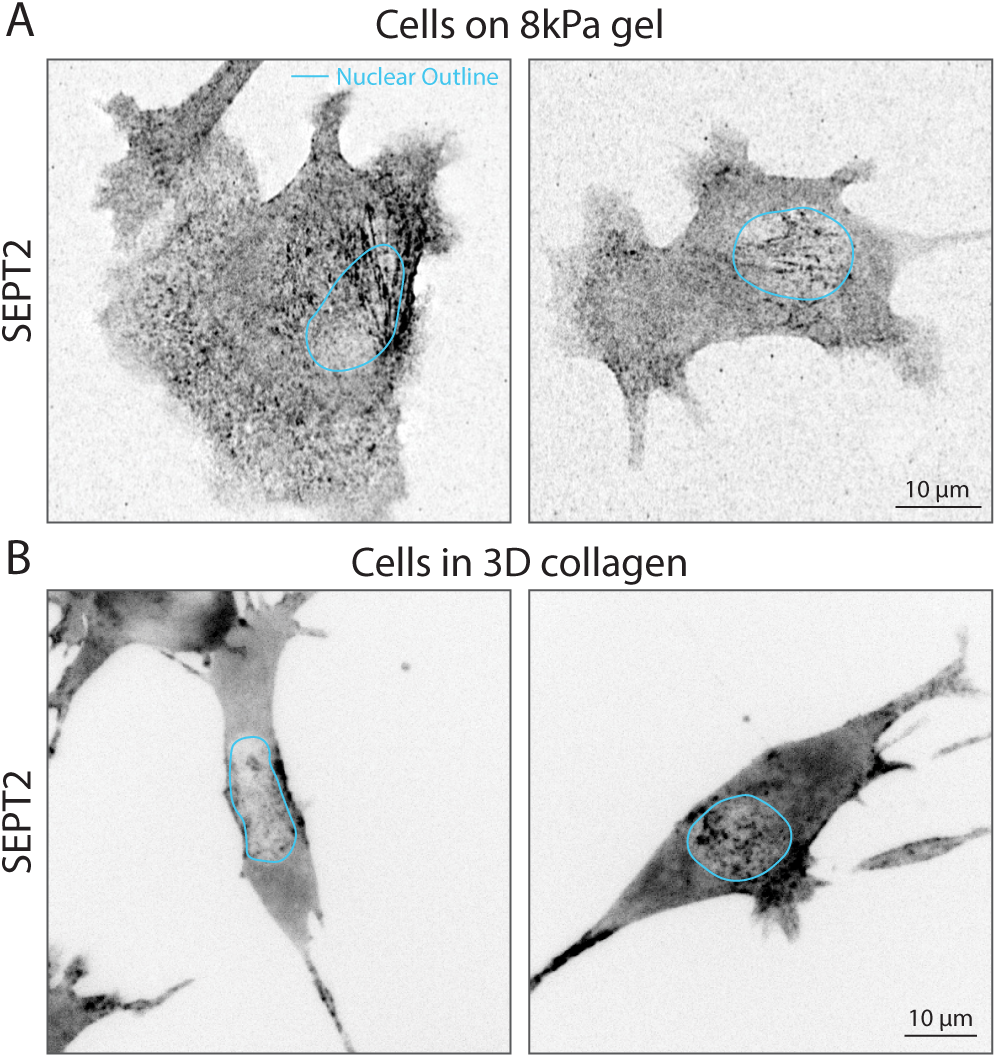
Septins colocalize with the nucleus in physiologic contexts. **A)** SEPT2-Halo cells labled with Halo dye embedded in a 3D collagen matrix. **B)** SEPT2-Halo cells labeled with Halo dye plated on polyacrylamide hydrogel with a shear modulus of 8 kPa. The nucleus is outlined in blue in each image.

**Figure S3.**
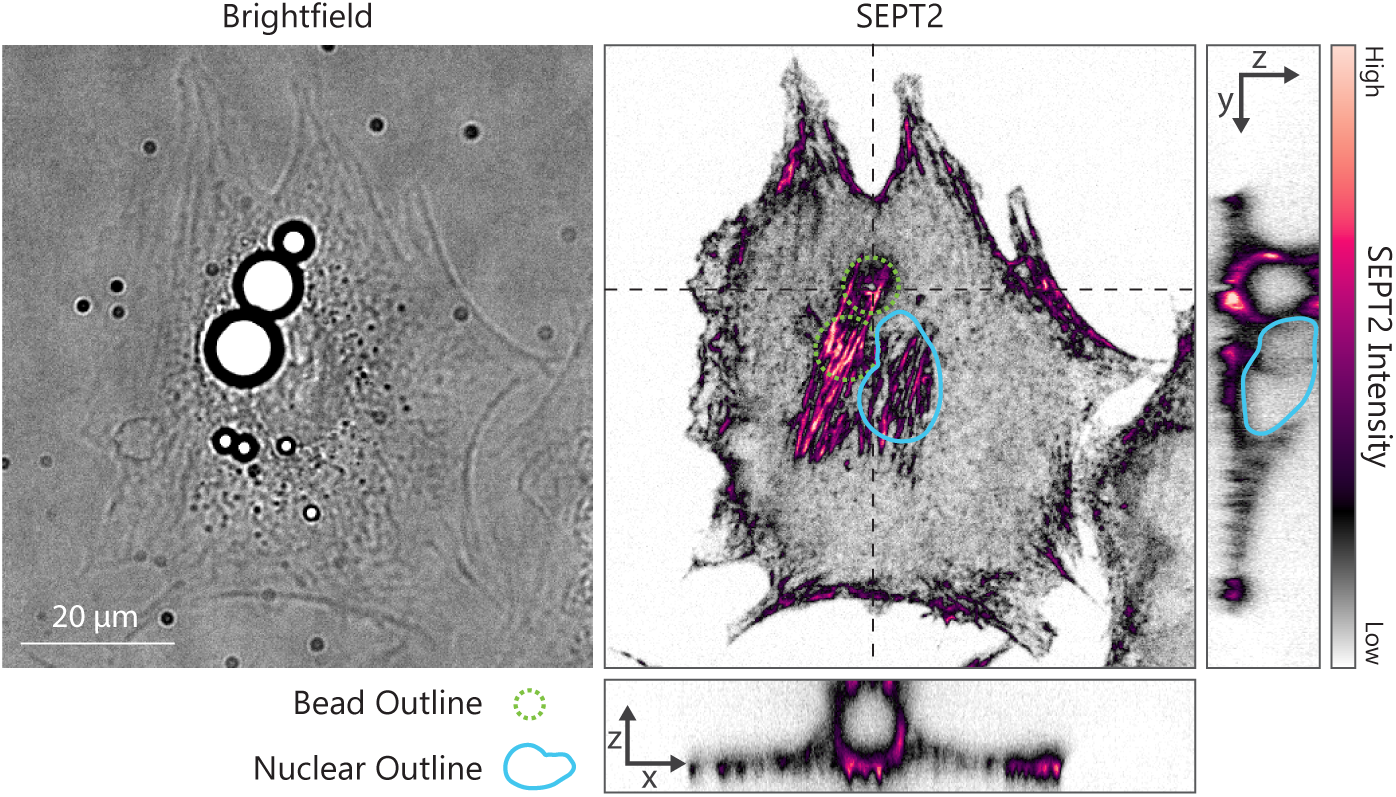
Glass beads induce septin filament assembly. A brightfield images shows two large beads internalized by the cell (and multiple smaller beads not internalized). The fluorescence image of the SEPT2-Halo cell shows septins accumulate under the large internalized beads (outlined in green dashed line). The nucleus is outlined in blue. X-Z and Y-Z projections show the bead fully internalized in the cell and septin accumulation under the bead.

**Figure S4.**
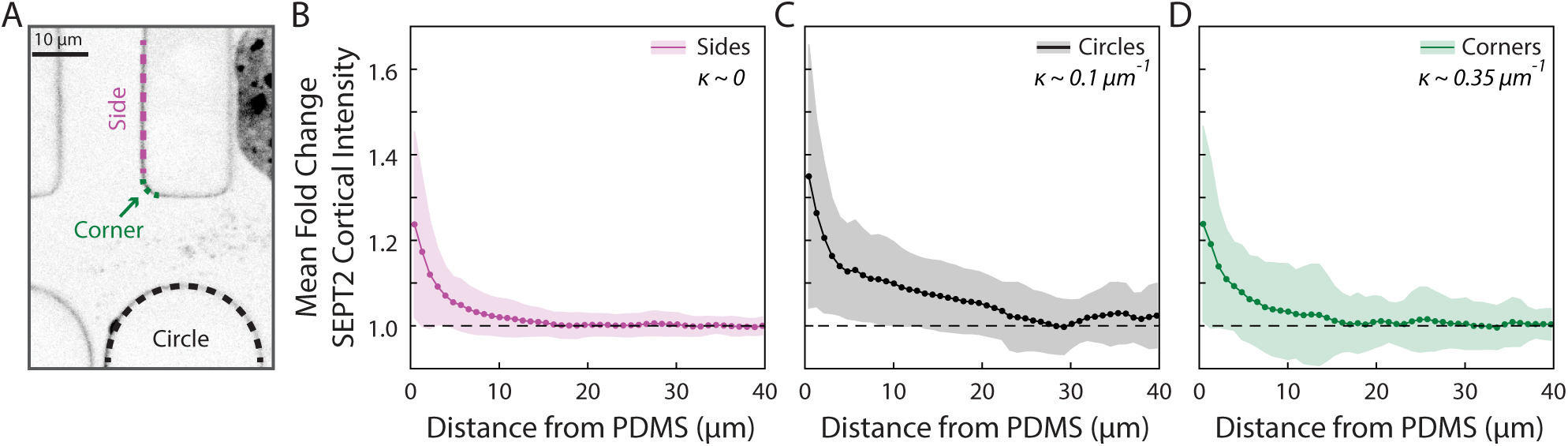
Cortical enrichment of septin does not depend on surface curvature. **A)** A representative image of the microfluidic device with labels identifying example surfaces with different curvature (*κ* = 1*/R*). **B-D)** Mean fold change in SEPT2 cortical assembly along flat sides (B; *κ* ∼ 0), circles (C; *κ* ∼ 0.1 *µm*^−1^), or corners (D; *κ* ∼ 0.35 *µm*^−1^) of features as a function of distance from the nucleus. Data for B-D represents the normalized average of all points along the cortex taken from 31 different cells, across 19 fields of view, from 3 different experiments. All data is normalized to the average SEPT2 intensity at the 30 *µ*m distance (dashed line), representing the average cortex intensity in the absence of the nucleus. Points represent the mean value and the shaded regions are the standard deviation.

**Figure S5.**
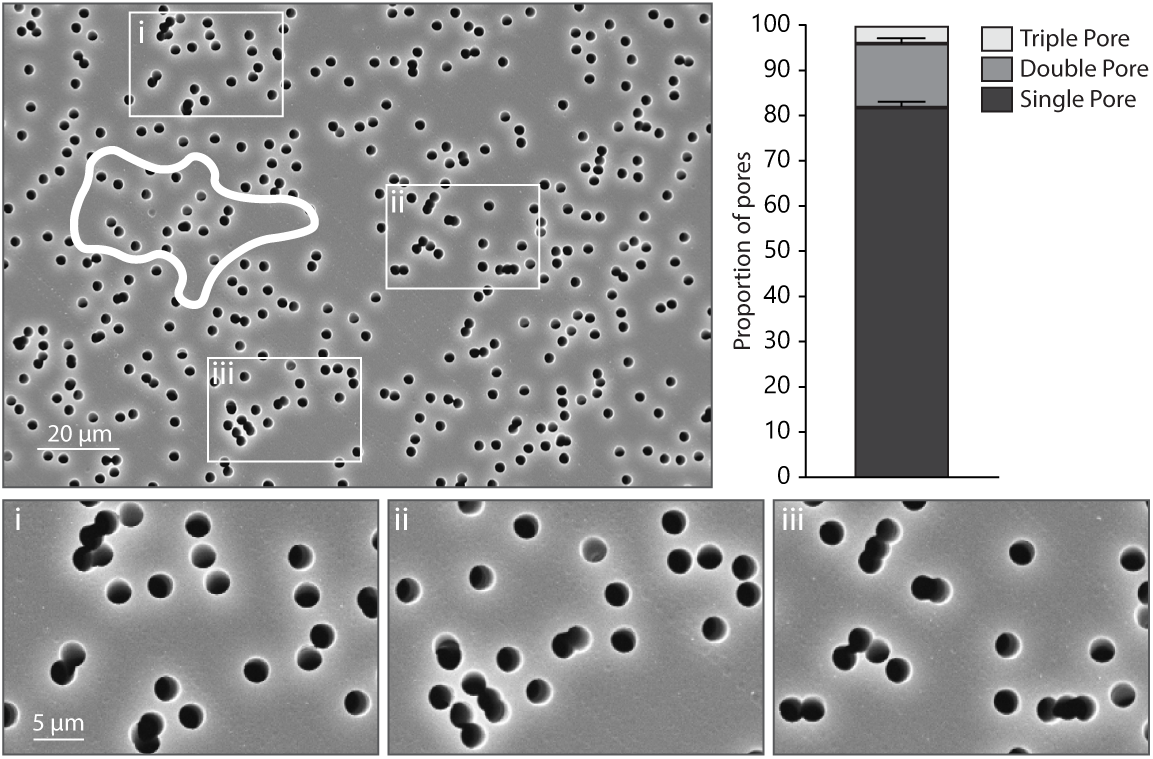
The distribution of pores in transwell plates are not uniform. An SEM image of a transwell insert showing non-uniform distribution of pores. Insets show multiple examples of pores overlapping each other, making pores significantly larger than the stated 3 *µ*m diameter, and possibly adding noise to the assay. The white solid line shows an outline of a representative cell on the transwell for scale. The bar plot shows a quantification of the pore size. Approximately 20% of pores are larger than 3 *µ*m in diameter.

### Supplemental Movies

**Movie M1: Septin networks coincide and move with the nucleus during migration.** A SEPT2-Halo cell stained with Hoechst nuclear dye migrating on a glass coverslip as shown in Fig. 1. The septin network under the nucleus is dynamically reorganized to remain under the nucleus as the cell migrates. The movie is 13 hours in length and is played at a frame rate of 10 fps. Time is in hr:min.

**Movie M2: Septin networks assemble under the nucleus in response to externally applied compression.** The ventral surface of a SEPT2-Halo cell is imaged prior to, during, and post 5 *µ*m confinement, which applies compression to the nucleus as shown in Fig. 3. Images were acquired every minute for 8 minutes (pre-confinement), 33 minutes (during confinement) and 15 minutes (post-confinement). The movie plays at 12 fps and time is in min:sec.

**Movie M3: Septin accumulates around the nucleus during confined migration.** A SEPT2-Halo cell migrating through a constriction in a microfluidic shows strong septin accumulation around the nucleus as it is squeezed between the pillars in the device as shown in Fig. 4A. The nucleus is stained with Hoescht dye. The movie is 6 hours in length and is played at a frame rate of 15 fps. Time is in hr:min.

**Movie M4: Septin accumulates where the nucleus pushes on the cortex.** A SEPT2-Halo cell migrating through a microfluidic device without being constricted still shows septin accumulation along the cortex as shown in Fig. 4C. The nucleus is stained with Hoescht dye. Movie is 1.75 hours in length and is played at a frame rate of 15 fps. Time is in hr:min.

## References

1. K. M. Yamada and M. Sixt. Mechanisms of 3D cell migration. Nat. Rev. Mol. Cell Biol., 100:1–15, October 2019. PMID: 31582855.

2. J. Lammerding. Mechanics of the nucleus. Comprehensive Physiology, 1(2):783–807, April 2011. PMID: 23737203.

3. C. M. Denais, R. M. Gilbert, P. Isermann, A. L. McGregor, M. te Lindert, B. Weigelin, P. M. Davidson, P. Friedl, K. Wolf, and J. Lammerding. Nuclear envelope rupture and repair during cancer cell migration. Science (New York, N.Y.), 352(6283):353–358, April 2016. PMID: 27013428.

4. T. P. Lele, R. B. Dickinson, and G. G. Gundersen. Mechanical principles of nuclear shaping and positioning. The Journal of Cell Biology, 217(10):3330–3342, October 2018. PMID: 30194270.

5. J. X. r. Renkawitz, A. Kopf, J. Stopp, I. Vries, M. K. Driscoll, J. Merrin, R. Hauschild, E. S. Welf, G. Danuser, R. Fiolka, and M. Sixt. Nuclear positioning facilitates amoeboid migration along the path of least resistance. Nature, 42:1–29, March 2019. PMID: 30944468.

6. A. J. Lomakin, C. J. Cattin, D. Cuvelier, Z. Alraies, M. Molina, G. P. F. Nader, N. Srivastava, P. J. Sáez, J. M. Garcia-Arcos, I. Y. Zhitnyak, A. Bhargava, M. K. Driscoll, E. S. Welf, R. Fiolka, R. J. Petrie, N. S. De Silva, J. M. González-Granado, N. Manel, A. M. Lennon-Duménil, D. J. Müller, and M. Piel. The nucleus acts as a ruler tailoring cell responses to spatial constraints. Science, 370(6514):eaba2894, October 2020. PMID: 33060332.

7. V. Venturini, F. Pezzano, F. C. Castro, H.-M. Häkkinen, S. Jiménez-Delgado, M. Colomer-Rosell, M. Marro, Q. Tolosa-Ramon, S. Paz-López, M. A. Valverde, J. Weghuber, P. Loza-Alvarez, M. Krieg, S. Wieser, and V. Ruprecht. The nucleus measures shape changes for cellular proprioception to control dynamic cell behavior. Science, 370(6514), October 2020. PMID: 33060331.

8. L. Lindenboim, H. Zohar, H. J. Worman, and R. Stein. The nuclear envelope: target and mediator of the apoptotic process. Cell Death Discovery, 6(1):29, April 2020. PMID: 32351716.

9. A. F. Pegoraro, P. Janmey, and D. A. Weitz. Mechanical Properties of the Cytoskeleton and Cells. Cold Spring Harb. Perspect. Biol., 9(11), November 2017.

10. G. F. Weber, M. A. Bjerke, and D. W. DeSimone. A mechanoresponsive cadherin-keratin complex directs polarized protrusive behavior and collective cell migration. Developmental Cell, 22(1):104–115, January 2012. PMID: 22169071.

11. O. Esue, A. A. Carson, Y. Tseng, and D. Wirtz. A Direct Interaction between Actin and Vimentin Filaments Mediated by the Tail Domain of Vimentin*. Journal of Biological Chemistry, 281(41):30393–30399, October 2006. PMID: 16901892.

12. V. Pelletier, N. Gal, P. Fournier, and M. L. Kilfoil. Microrheology of Microtubule Solutions and Actin-Microtubule Composite Networks. Physical Review Letters, 102(18):188303, May 2009. PMID: 19518917.

13. L. Farhadi, S. N. Ricketts, M. J. Rust, M. Das, R. M. Robertson-Anderson, and J. L. Ross. Actin and microtubule crosslinkers tune mobility and control co-localization in a composite cytoskeletal network. Soft Matter, 16(31):7191–7201, August 2020. PMID: 32207504.

14. J. P. Conboy, M. G. Lettinga, P. Boukany, F. C. MacKintosh, and G. H. Koenderink. Actin and vimentin jointly control cell viscoelasticity and compression stiffening. Mol. Biol. Cell, January 2026. PMID: 41400926.

15. M. Kinoshita, C. M. Field, M. L. Coughlin, A. F. Straight, and T. J. Mitchison. Self- and actin-templated assembly of Mammalian septins. Dev. Cell, 3(6):791–802, December 2002.

16. B. E. Kremer, L. A. Adang, and I. G. Macara. Septins Regulate Actin Organization and Cell Cycle Arrest Through SOCS7-Mediated Nuclear Accumulation of NCK. Cell, 130(5):837–850, September 2007. PMID: 17803907.

17. F. Calvo, R. Ranftl, S. Hooper, A. J. Farrugia, E. Moeendarbary, A. Bruckbauer, F. Batista, G. Charras, and E. Sahai. Cdc42EP3/BORG2 and Septin Network Enables Mechano-transduction and the Emergence of Cancer-Associated Fibroblasts. Cell Rep., 13(12):2699–2714, December 2015. PMID: 26711338.

18. C. S. Martins, C. Taveneau, G. Castro-Linares, M. Baibakov, N. Buzhinsky, M. Eroles, V. Milanović, S. Omi, J.-D. Pedelacq, F. Iv, L. Bouillard, A. Llewellyn, M. Gomes, M. Belhabib, M. Kuzmić, P. Verdier-Pinard, S. Lee, A. Badache, S. Kumar, C. Chandre, S. Brasselet, F. Rico, O. Rossier, G. H. Koenderink, J. Wenger, S. Cabantous, and M. Mavrakis. Human septins organize as octamer-based filaments and mediate actin-membrane anchoring in cells. J. Cell Biol., 222(3), March 2023. PMID: 36562751.

19. W. Sturgess, S. Packirisamy, R. Geneidy, P. Nordenfelt, and V. Swaminathan. ECM-dependent regulation of septin 7 in focal adhesions promotes mechanosensing and functional response in fibroblasts. iScience, 27(12):111355, December 2024. PMID: 39650732.

20. I. A. Cavini, D. A. Leonardo, H. V. D. Rosa, D. K. S. V. Castro, H. D’Muniz Pereira, N. F. Valadares, A. P. U. Araujo, and R. C. Garratt. The Structural Biology of Septins and Their Filaments: An Update. Front Cell Dev Biol, 9:765085, November 2021. PMID: 34869357.

21. X. Bai, J. R. Bowen, T. K. Knox, K. Zhou, M. Pendziwiat, G. Kuhlenbäumer, C. V. Sindelar, and E. T. Spiliotis. Novel septin 9 repeat motifs altered in neuralgic amyotrophy bind and bundle microtubules. Journal of Cell Biology, 203(6):895–905, December 2013. PMID: 24344182.

22. M. Mavrakis, Y. Azou-Gros, F.-C. Tsai, J. Alvarado, A. Bertin, F. Iv, A. Kress, S. Brasselet, G. H. Koenderink, and T. Lecuit. Septins promote F-actin ring formation by crosslinking actin filaments into curved bundles. Nat. Cell Biol., 16(4):322–334, April 2014. PMID: 24633326.

23. E. T. Spiliotis. Spatial effects - site-specific regulation of actin and microtubule organization by septin GTPases. J. Cell Sci., 131(1), January 2018. PMID: 29326311.

24. M. Ihara, A. Kinoshita, S. Yamada, H. Tanaka, A. Tanigaki, A. Kitano, M. Goto, K. Okubo, H. Nishiyama, O. Ogawa, C. Takahashi, S. Itohara, Y. Nishimune, M. Noda, and M. Kinoshita. Cortical organization by the septin cytoskeleton is essential for structural and mechanical integrity of mammalian spermatozoa. Developmental Cell, 8(3):343–352, March 2005. PMID: 15737930.

25. S. Mostowy and P. Cossart. Septins: the fourth component of the cytoskeleton. Nat. Rev. Mol. Cell Biol., 13(3):183–194, February 2012. PMID: 22314400.

26. M. Lam and F. Calvo. Regulation of mechanotransduction: Emerging roles for septins. Cytoskeleton, 76(1):115–122, September 2018. PMID: 30091182.

27. M. L. Berre, J. Aubertin, and M. Piel. Fine control of nuclear confinement identifies a threshold deformation leading to lamina rupture and induction of specific genes. Integr. Biol., 4(11):1406–1409, November 2012. PMID: 23038068.

28. J. Keys, A. Windsor, and J. Lammerding. Assembly and Use of a Microfluidic Device to Study Cell Migration in Confined Environments. Methods in Molecular Biology (Clifton, N.J.), 1840:101–118, August 2018. PMID: 30141042.

29. A. A. Bridges, M. S. Jentzsch, P. W. Oakes, P. Occhipinti, and A. S. Gladfelter. Micron-scale plasma membrane curvature is recognized by the septin cytoskeleton. J. Cell Biol., 213(1):23–32, April 2016. PMID: 27044896.

30. K. S. Cannon, B. L. Woods, J. M. Crutchley, and A. S. Gladfelter. An amphipathic helix enables septins to sense micrometer-scale membrane curvature. J. Cell Biol., pages jcb.201807211–10, January 2019. PMID: 30659102.

31. K. Nakazawa, G. Kumar, B. Chauvin, A. Di Cicco, L. Pellegrino, M. Trichet, B. Hajj, J. Cabral, A. Sain, S. Mangenot, and A. Bertin. A human septin octamer complex sensitive to membrane curvature drives membrane deformation with a specific mesh-like organization. Journal of Cell Science, 136(11):jcs260813, June 2023. PMID: 37305997.

32. D. M. Graham, T. Andersen, L. Sharek, G. Uzer, K. Rothenberg, B. D. Hoffman, J. Rubin, M. Balland, J. E. Bear, and K. Burridge. Enucleated cells reveal differential roles of the nucleus in cell migration, polarity, and mechanotransduction. J. Cell Biol., 217(3):895–914, March 2018. PMID: 29351995.

33. S. Marcotti, D. B. de Freitas, L. D. Troughton, F. N. Kenny, T. J. Shaw, B. M. Stramer, and P. W. Oakes. A workflow for rapid unbiased quantification of fibrillar feature alignment in biological images. Front Comput Sci, 3, October 2021. PMID: 34888522.

34. A. J. Tooley, J. Gilden, J. Jacobelli, P. Beemiller, W. S. Trimble, M. Kinoshita, and M. F. Krummel. Amoeboid T lymphocytes require the septin cytoskeleton for cortical integrity and persistent motility. Nat. Cell Biol., 11(1):17–26, January 2009. PMID: 19043408.

35. J. Gilden and M. F. Krummel. Control of cortical rigidity by the cytoskeleton: emerging roles for septins. Cytoskeleton, 67(8):477–486, August 2010. PMID: 20540086.

36. S. Mostowy, S. Janel, C. Forestier, C. Roduit, S. Kasas, J. Pizarro-Cerdá, P. Cossart, and F. Lafont. A role for septins in the interaction between the Listeria monocytogenes INVASION PROTEIN InlB and the Met receptor. Biophys. J., 100(8):1949–1959, April 2011. PMID: 21504731.

37. T. J. Park, S. K. Kim, and J. B. Wallingford. The planar cell polarity effector protein Wdpcp (Fritz) controls epithelial cell cortex dynamics via septins and actomyosin. Biochem. Biophys. Res. Commun., 456(2):562–566, January 2015. PMID: 25436430.

38. J. Okletey, D. Angelis, T. M. Jones, C. Montagna, and E. T. Spiliotis. An oncogenic isoform of septin 9 promotes the formation of juxtanuclear invadopodia by reducing nuclear deformability. Cell Reports, 42(8):112893, July 2023. PMID: 37516960.

39. Q. Hu, L. Milenkovic, H. Jin, M. P. Scott, M. V. Nachury, E. T. Spiliotis, and W. J. Nelson. A septin diffusion barrier at the base of the primary cilium maintains ciliary membrane protein distribution. Science, 329(5990):436–439, July 2010. PMID: 20558667.

40. H. Ewers, T. Tada, J. D. Petersen, B. Racz, M. Sheng, and D. Choquet. A Septin-Dependent Diffusion Barrier at Dendritic Spine Necks. PLoS One, 9(12):e113916, December 2014. PMID: 25494357.

41. J. K. Gilden, S. Peck, Y.-C. M. Chen, and M. F. Krummel. The septin cytoskeleton facilitates membrane retraction during motility and blebbing. J. Cell Biol., 196(1):103–114, January 2012. PMID: 22232702.

42. N. Vadnjal, S. Nourreddine, G. Lavoie, M. Serres, P. P. Roux, and E. K. Paluch. Proteomic analysis of the actin cortex in interphase and mitosis. J. Cell Sci., July 2022. PMID: 35892282.

43. A. D. Weems, E. S. Welf, M. K. Driscoll, F. Y. Zhou, H. Mazloom-Farsibaf, B.-J. Chang, V. S. Murali, G. M. Gihana, B. G. Weiss, J. Chi, D. Rajendran, K. M. Dean, R. Fiolka, and G. Danuser. Blebs promote cell survival by assembling oncogenic signalling hubs. Nature, 615(7952):517–525, March 2023. PMID: 36859545.

44. V. Stjepić, M. Nakamura, J. Hui, and S. M. Parkhurst. Two Septin complexes mediate actin dynamics during cell wound repair. Cell Reports, 43(5), May 2024. PMID: 38728140.

45. H. Herrmann and U. Aebi. Intermediate Filaments: Structure and Assembly. Cold Spring Harbor Perspectives in Biology, 8(11):a018242, November 2016. PMID: 27803112.

46. P. A. Coulombe. Discovery of keratin function and role in genetic diseases: the year that 1991 was. Molecular Biology of the Cell, 27(18):2807–2810, September 2016. PMID: 27634744.

47. A. E. Patteson, A. Vahabikashi, K. Pogoda, S. A. Adam, K. Mandal, M. Kittisopikul, S. Sivagurunathan, A. Goldman, R. D. Goldman, and P. A. Janmey. Vimentin protects cells against nuclear rupture and DNA damage during migration. J. Cell Biol., 218(12):4079–4092, December 2019. PMID: 31676718.

48. I. Dupin, Y. Sakamoto, and S. Etienne-Manneville. Cytoplasmic intermediate filaments mediate actin-driven positioning of the nucleus. Journal of Cell Science, 124(Pt 6):865–872, March 2011. PMID: 21378307.

49. G. Joberty, R. R. Perlungher, P. J. Sheffield, M. Kinoshita, M. Noda, T. Haystead, and I. G. Macara. Borg proteins control septin organization and are negatively regulated by Cdc42. Nat. Cell Biol., 3(10):861–866, October 2001. PMID: 11584266.

50. S. Chandrasekar, M. E. Utgaard, B. Somerfield, H. Wu, J. R. Beach, and P. W. Oakes. Local RhoA activation induces septin recruitment, September 2025. ISSN: 2692-8205 Pages: 2025.09.29.679255 Section: New Results.

51. M. Kinoshita. The septins. Genome Biol., 4(11):236, October 2003. PMID: 14611653.

52. E. P. Karasmanis, D. Hwang, K. Nakos, J. R. Bowen, D. Angelis, and E. T. Spiliotis. A Septin Double Ring Controls the Spatiotemporal Organization of the ESCRT Machinery in Cytokinetic Abscission. Current Biology, 29(13):2174–2182.e7, July 2019. PMID: 31204162.

53. A. Bertin, M. A. McMurray, L. Thai, G. Garcia, V. Votin, P. Grob, T. Allyn, J. Thorner, and E. Nogales. Phosphatidylinositol-4,5-bisphosphate promotes budding yeast septin filament assembly and organization. Journal of Molecular Biology, 404(4):711–731, December 2010. PMID: 20951708.

54. M. Omrane, A. S. Camara, C. Taveneau, N. Benzoubir, T. Tubiana, J. Yu, R. Guérois, D. Samuel, B. Goud, C. Poüs, S. Bressanelli, R. C. Garratt, A. R. Thiam, and A. Gassama-Diagne. Septin 9 has Two Polybasic Domains Critical to Septin Filament Assembly and Golgi Integrity. iScience, 13:138–153, March 2019. PMID: 30831549.

55. J. J. Lacroix and T. D. Wijerathne. PIEZO channels as multimodal mechanotransducers. Biochemical Society Transactions, 53(1):293–302, February 2025. PMID: 39936392.

56. G. W. G. Luxton, E. R. Gomes, E. S. Folker, E. Vintinner, and G. G. Gundersen. Linear arrays of nuclear envelope proteins harness retrograde actin flow for nuclear movement. Science, 329(5994):956–959, August 2010. PMID: 20724637.

57. S. Antoku, R. Zhu, S. Kutscheidt, O. T. Fackler, and G. G. Gundersen. Reinforcing the LINC complex connection to actin filaments: the role of FHOD1 in TAN line formation and nuclear movement. Cell Cycle, 14(14):2200–2205, June 2015. PMID: 26083340.

58. F. Caudron and Y. Barral. Septins and the lateral compartmentalization of eukaryotic membranes. Developmental Cell, 16(4):493–506, April 2009. PMID: 19386259.

59. A. S. Zhovmer, A. Manning, C. Smith, A. Nguyen, O. Prince, P. J. Sáez, X. Ma, D. Tsygankov, A. X. Cartagena-Rivera, N. A. Singh, R. K. Singh, and E. D. Tabdanov. Septins provide microenvironment sensing and cortical actomyosin partitioning in motile amoeboid T lymphocytes. Science Advances, 10(1):eadi1788, January 2024. PMID: 38170778.

60. M. Scott, W. G. McCluggage, K. J. Hillan, P. A. Hall, and S. E. H. Russell. Altered patterns of transcription of the septin gene, SEPT9, in ovarian tumorigenesis. Int. J. Cancer, 118(5):1325–1329, March 2006. PMID: 16161048.

61. L. Stanbery, N. J. D’Silva, J. S. Lee, C. R. Bradford, T. E. Carey, M. E. Prince, G. T. Wolf, F. P. Worden, K. G. Cordell, and E. M. Petty. High SEPT9_v1 Expression Is Associated with Poor Clinical Outcomes in Head and Neck Squamous Cell Carcinoma. Translational Oncology, 3(4):239–245, August 2010. PMID: 20689765.

62. R. Gilad, K. Meir, I. Stein, L. German, E. Pikarsky, and N. J. Mabjeesh. High SEPT9_i1 protein expression is associated with high-grade prostate cancers. PloS One, 10(4):e0124251, 2015. PMID: 25898316.

63. P. Jȩdrzejczak, K. Saramowicz, J. Kuś, J. Barczuk, W. Rozpȩdek-Kaminśka, N. Siwecka, G. Galita, W. Wiese, and I. Majsterek. SEPT9_i1 and Septin Dynamics in Oncogenesis and Cancer Treatment. Biomolecules, 14(9):1194, September 2024. PMID: 39334960.

64. F. Mayca-Pozo, S. M. Butts, A. W. Schaefer, and E. T. Spiliotis. Septins buffer actomyosin forces to protect the nucleus from genotoxic mechanical stress. *Submitted*, January 2026.

65. P. Krausewitz, N. Kluemper, A.-P. Richter, T. Büttner, G. Kristiansen, M. Ritter, and J. Ellinger. Early Dynamics of Quantitative SEPT9 and SHOX2 Methylation in Circulating Cell-Free Plasma DNA during Prostate Biopsy for Prostate Cancer Diagnosis. Cancers, 14(18):4355, September 2022. PMID: 36139516.

66. L. de Vos, M. Jung, R.-M. Koerber, E. G. Bawden, T. A. W. Holderried, J. Dietrich, F. Bootz, P. Brossart, G. Kristiansen, and D. Dietrich. Treatment Response Monitoring in Patients with Advanced Malignancies Using Cell-Free SHOX2 and SEPT9 DNA Methylation in Blood: An Observational Prospective Study. The Journal of molecular diagnostics: JMD, 22(7):920–933, July 2020. PMID: 32361006.

67. S.-L. Zhang, H.-J. Yu, Z.-Q. Lian, J. Wan, S.-M. Xie, W. Lei, Q.-P. Chen, L. Zhang, and Q. Wang. Septin9 DNA methylation is associated with breast cancer recurrence or metastasis. The Journal of International Medical Research, 52(1):3000605231220827, January 2024. PMID: 38180895.

68. H. W. Johnson and M. J. Schell. Neuronal IP3 3-kinase is an F-actin-bundling protein: role in dendritic targeting and regulation of spine morphology. Molecular Biology of the Cell, 20(24):5166–5180, December 2009. PMID: 19846664.

69. F. A. Ran, P. D. Hsu, J. Wright, V. Agarwala, D. A. Scott, and F. Zhang. Genome engineering using the CRISPR-Cas9 system. Nature Protocols, 8(11):2281–2308, November 2013. PMID: 24157548.

70. A. D. Doyle. Generation of 3D collagen gels with controlled, diverse architectures. Current protocols in cell biology / editorial board, Juan S. Bonifacino … [et al.], 72:10.20.1–10.20.16, September 2016. PMID: 27580704.

71. P. W. Oakes, T. C. Bidone, Y. Beckham, A. V. Skeeters, G. R. R.-S. Juan, S. P. Winter, G. A. Voth, and M. L. Gardel. Lamellipodium is a myosin-independent mechanosensor. Proc. Natl. Acad. Sci. U. S. A., 115(11):2646–2651, March 2018. PMID: 29487208.

72. R. Kumari, K. Ven, M. Chastney, S. B. Kokate, J. Peränen, J. Aaron, K. Kogan, L. Almeida-Souza, E. Kremneva, R. Poincloux, T.-L. Chew, P. W. Gunning, J. Ivaska, and P. Lappalainen. Focal adhesions contain three specialized actin nanoscale layers. Nature Communications, 15(1):2547, March 2024. PMID: 38514695.

73. A. Stubb, C. Guzmán, E. Närvä, J. Aaron, T.-L. Chew, M. Saari, M. Miihkinen, G. Jacquemet, and J. Ivaska. Superresolution architecture of cornerstone focal adhesions in human pluripotent stem cells. Nature Communications, 10(1):4756, October 2019. PMID: 31628312.

74. P. Kanchanawong, G. Shtengel, A. M. Pasapera, E. B. Ramko, M. W. Davidson, H. F. Hess, and C. M. Waterman. Nanoscale architecture of integrin-based cell adhesions. Nature, 468(7323):580–584, November 2010. PMID: 21107430.

75. L. B. Case, M. A. Baird, G. Shtengel, S. L. Campbell, H. F. Hess, M. W. Davidson, and C. M. Waterman. Molecular mechanism of vinculin activation and nanoscale spatial organization in focal adhesions. Nat. Cell Biol., June 2015. PMID: 26053221.

76. G. Shtengel, J. A. Galbraith, C. G. Galbraith, J. Lippincott-Schwartz, J. M. Gillette, S. Manley, R. Sougrat, C. M. Waterman, P. Kanchanawong, M. W. Davidson, R. D. Fetter, and H. F. Hess. Interferometric fluorescent super-resolution microscopy resolves 3D cellular ultrastructure. Proc. Natl. Acad. Sci. U. S. A., 106(9):3125–3130, March 2009. PMID: 19202073.

77. J. Keys, B. C. H. Cheung, M. A. Elpers, M. Wu, and J. Lammerding. Rear cortex contraction aids in nuclear transit during confined migration by increasing pressure in the cell posterior. Journal of Cell Science, 137(12):jcs260623, June 2024. PMID: 38832512.

